# Targeted inhibition of Bcl-xL following radiation reduces tumourigenesis in preclinical models of H3K27M-altered diffuse midline glioma

**DOI:** 10.1101/2024.12.20.629607

**Authors:** Ashley Vardon, Scott Haston, Romain Guiho, Diana Carvahlo, Rebecca Carter, Jessica Boult, Daniel Gharai, Diwakar Santhakumar, Ielyaas Cloete, James Gray, John Apps, Olumide Ogunbiyi, Daohong Zhou, Guangrong Zheng, Ramón Martínez-Máñez, Mark Lythgoe, Laura Donovan, Angel Montero Carcaboso, Owen Williams, Jesus Gil, Thomas Jacques, Jamie Dean, David Michod, Chris Jones, Darren Hargrave, Juan-Pedro Martinez-Barbera

**Author notes:** Corresponding author: Prof. JP Martinez-Barbera, Developmental Biology and Cancer Programme, Birth Defects Research Centre, Great Ormond Street Institute of Child Health, University College London, London, UK.

## Abstract

**Background:** Diffuse midline gliomas (DMGs) with histone H3K27M mutations represent a devastating paediatric brain cancer characterized by abysmal prognosis and limited treatment options. The only approved treatment is radiotherapy (RT), but most of the tumours relapse with fatal consequences. In this study, we sought to investigate whether irradiation leads to senescence induction and explore the efficacy of senolytics against DMG.

**Methods:** We have characterised the senescent phenotype of five genetically heterogeneous H3K27M-altered human DMG cell lines, combining cellular and/or molecular approaches. The sensitivity of senescent cells to Bcl-xL inhibition has been demonstrated in dose/response curves in vitro and in a PDX model of DMG.

**Results:** Here, we show that ionizing radiation induces senescence and SASP responses in both TP53 mutant and wild-type H3K27M-altered human DMG cell lines. We identify Navitoclax as a potent senolytic agent that selectively targets senescent DMG cells into apoptosis by inhibiting Bcl-xL. Related compounds, such as a proteolysis-targeting chimera (PROTAC)-mediated Bcl-xL degradation and a galacto-conjugated form of Navitoclax also show an effective senolytic activity in senescent cancer cells. Finally, we show that a combination therapy of irradiation and Navitoclax results in reduced tumor burden and increased mouse survival in an orthotopic xenograft DMG model.

**Conclusion:** These results offer a rationale for further clinical development of senolytic therapies as part of multimodal treatment approaches for DMG patients/

**Key Points:**

- Ionising irradiation induces senescence in human DMG cells independently of the p53 status.
- Bcl-xL inhibition results in apoptosis of human DMG senescent cells in synergy with irradiation.
- Combination of irradiation and BcL-xL inhibition reduces tumourigenesis in a PDX model of DMG.

**Importance of the Study:** H3K27M-altered DMG are devastating paediatric tumours with an abysmal prognosis. The only approved treatment is radiotherapy but this is palliative and tumours almost always relapse with fatal consequences for the patients. In this study, we show that radiotherapy results in senescence induction in five genomically heterogeneous human DMG cell lines. We identify that drugs targeting the anti-apoptotic protein Bcl-xL show a strong senolytic activity in conjunction with radiotherapy both in vitro in DMG cells and in vivo in a PDX model of H3K27M-altered DMG. Treatment with Bcl-xL inhibitor Navitoclax, or related compounds targeting Bcl-xL protein degradation or containing a galactose conjugated form of Navitoclax results in DMG cancer cell apoptosis. As several of these inhibitors are currently being tested in ongoing clinical trials against other diseases, our data support the use of Bcl-xL inhibition mediated senolytics as an adjuvant therapy to radiotherapy to potentially improve outcomes in this challenging disease setting.

## Introduction

H3K27M-altered diffuse midline glioma (DMG) of the pons represents around 10% of childhood brain tumours, and has a peak incidence of 6-8 years. Despite four decades of clinical trials, the only proven effective therapy is radiotherapy (RT), and unfortunately this is palliative and most patients relapse and succumb to the diseases with a median survival of 8-11 months post-diagnosis ^1^. RT can induce cellular senescence in a variety of cancer contexts including brain tumours^2–5^. Lingering senescent cells within the irradiated tumour bed may contribute to create a permissive environment that promote relapse. Therefore, targeting of these senescent cells may represent a novel therapeutic strategy for combating DMG.

Cellular senescence is characterised by stable cell cycle arrest, which is maintained by critical pathways regulating cell cycle progression (e.g. p53/p21 and p16/RB)^6^. Senescent cells can elicit cell non-autonomous activities through the Senescence-Associated Secretory Phenotype (SASP), a complex secretory programme composed of a multitude of cytokines and chemokines (e.g. IL1α, IL1β, IL6), growth factors (e.g. EGF, FGFs, VEGF), and other active chemicals^7^. Although transient SASP activation has beneficial physiological effects, such as wound healing^8^, chronic SASP activation can promote proliferation of transformed cells and/or create a permissive microenvironment that supports tumour progression, malignancy and metastasis^9^. Of clinical relevance, senescent cell ablation or modulation of the SASP, can reduce tumour burden, increase mouse survival, decrease tumour relapse and alleviate the negative effects of anticancer treatment^10–12^.

Senescent cells can be ablated effectively using senolytic compounds, which exploit molecular vulnerabilities in senescent cells Ovadya and Krizhanovsky 2018). Several drugs that can selectively kill senescent cells have been identified, including dasatinib and quercetin (referred as D+Q)^13^, Bcl2 family inhibitors such as ABT-263 (also known as Navitoclax) and ABT-737^14,15^, a modified FOXO4-p53 interfering peptide^16^, HSP90 inhibitors (e.g., alvespimycin)^17^, piperlongumine^18^, cardiac glycosides^19,20^, N-myristoylation inhibitors^21^ and β-galactosidase-activated nanoparticles and pro-drugs^22–24^. Among all senolytics, inhibition of Bcl-xL/Bcl-2, members of the anti-apoptotic Bcl-2 protein family, with BH3-mimetics is one of the most commonly used method to eliminate senescent cells pharmacologically both *in vitro* and *in vivo* across a variety of cell types ^25–31^.

In this study, we have characterised the effects of radiation regimens on a variety of molecularly distinct H3K27M-altered human DMG cellular models, demonstrating the induction of a senescence programme and activation of a SASP. We have identified Bcl-xL inhibition as a vulnerability of RT-induced senescent DMG cells and shown the preclinical efficacy of a set of BH3-mimetics, including several clinically approved drugs and phase I/II compounds.

## Methods

### Primary DMG cell line culture

H3K27M-altered human DMG cells (**Supplementary Table 1**) were grown under adherent stem cell conditions using flasks coated with laminin (10ug/ml) (R&D systems, Bio-techne) in tumour stem media complete (TSM-C). TSM base (TSM-B) was made using 250mL Neurobasal-A Medium (Invitrogen), 250mL DMEM/F-12 (Invitrogen), 1% HEPES buffer solution (ThermoFisher), Sodium pyruvate MEM 100mM (ThermoFisher), MEM non-essential amino acids solution 10mM (ThermoFisher), GlutaMAX-I supplement (ThermoFisher), Antibiotic-antimycotic (ThermoFisher). Prior to use TSM-B was supplemented with B-27 without vitamin A (Invitrogen) and the growth factors epidermal growth factor (EGF, 20 ng/ml Peprotech), basal fibroblast growth factor (bFGF, 20 ng/ml, Peprotech), platelet-derived growth factor (PDGF-AA and PDGF-BB, 10 ng/ml, Peprotech) and 0.2% heparin sulphate (Stemcell technologies), TSM-C was used within two weeks. All cell lines were routinely confirmed mycoplasma negative using the MycoAlert™ PLUS Mycoplasma Detection Kit (Lonza, catalogue no. LT07-710).

### RNA Sequencing and bioinformatic analysis

RNA was extracted from both non-irradiated (proliferative) and irradiated (senescent) human DMG cells using the Rneasy Micro kit (Qiagen). Total RNA integrity was validated using Agilent’s 4200 Tapestation, with all samples having RIN values >9.4. UCL genomics core facility prepared cDNA libraries using 250ng of total RNA and the KAPA mRNA HyperPrep Kit. High-quality libraries were confirmed on the Agilent TapeStation 4200. Samples were pooled, normalized to 4nM using the Normalase assay, and sequenced on the NextSeq 500 instrument with a 75bp single read run. Differential Expression Analysis with DESeq2 DESeq2 was employed for differential expression (DE) analysis, applying normalisation factors based on a negative binomial model. Independent filtering was used to include only genes with sufficient reads for differential expression detection. Gene ontology enrichment analysis considered the heteroskedastic nature of RNA sequencing data, utilizing the GOSeq package in R. Visualization of results was done using R. Gene Set Enrichment Analysis GSEA assessed ranked DESeq2 results using the Wald Statistic. Pre-ranked gene lists were obtained from MSigDB, and GSEA was performed with version 4.2.3 from the Broad Institute. The False Discovery Rate (FDR) was used to estimate the probability of false positive findings.

### Gamma-ray irradiation of DMG cells

137Cs γ-ray irradiation at a dose rate of 1.76 Gy/min was conducted using an IBL 437C (CIS Bio-International, Codolet, France). Primary DMG cells were cultured to less than passage 10 prior to induction of senescence through irradiation. Cells were Accutased (Sigma) counted using Trypan blue and cells resuspended in 1ml of TSM-C and irradiated in 15ml falcon tubes at doses described (He et al., 2011). To avoid false-positivity due to differences in confluency, irradiated (senescent) and proliferative cells were collected for molecular or cellular analyses at 70-80% confluency.

### Senescence-associated beta-galactosidase staining

Senescence was verified by staining for senescence-associated-β-galactosidase (SA-β-gal) using a commercially available SA-β-gal staining kit (Sigma, catalogue no. CS0030). Following manufacture instructions, as previously described (Gonzalez-Meljem, Apps et al. 2018), a minimum of 400 cells counted per condition. Positive (blue) cells are expressed as a percentage of total cell number.

### EdU staining

EdU (10µmol final concentration) diluted in TSM-C was added to the cells overnight. Click-it EdU (ThermoFisher) reaction was carried out as per manufacturers protocol. Briefly, media was removed, cells were washed with 1X PBS, and fixed with 4% Paraformaldehyde for 15 minutes. Cells were then washed with 3% BSA and permeabilised with 0.5% Triton-X100. Cells were washed a twice with 3% BSA and Click-it EdU reaction cocktail (Alexa Flour 488, 597 were used) was added to the cells for 30 minutes. A further wash with 3% BSA and nuclei were counter stained with Hoescht for 30 minutes.

### Drugs for in vitro and in vivo testing

Navitoclax (ABT-263, catalogue no. 11500, Cayman Chemical), ABT-737 (catalogue no. ab141336, Abcam), Dasatinib (catalogue no. S1021, Selleckchem), Quercetin (catalogue no. S2391, Selleckchem), Piperlongumine (catalogue no. CAY-11006, Selleckchem), Alvespimycin (catalogue no. hy-12024, Selleckchem), BCL specific agents A1155463 (catalogue no. S7800, Selleckchem), A1331852 (catalogue no. S7801, Selleckchem), A1210477 (catalogue no. S7790, Selleckchem), Obatoclax Mesylate (GX15-070) (catalogue no. S1057, Selleckchem). PROTACs DT2216 and PZ18753B were obtained from Daohong Zhou (University of Florida) under MTA (5934207). Nav-Gal was obtained from Ramón Martínez-Máñez (IDM, University of Valencia) (**Supplementary Table 2**). All drugs were dissolved in DMSO and stored in aliquots at −20°C. Drugs were diluted in cell culture medium and added to the cells for the durations and concentrations indicated. ABT-263 (100 mg, ApexBio), was weighed and vehicle (ethanol:polyethylene glycol 400:Phosal 50 PG) was added, individual aliquots were prepared, prior to administration drug was sonicated on ice into solution using Diagenode Bioruptor®, high frequency, 30 second intervals, for 15 minutes. ABT-263 was administered to mice by gavage at 50 mg per kg body weight per day (mg/kg/d) for 5 days for two cycles.

### Immunofluorescence staining

Immunofluorescence staining was carried out by first fixing wells of 96 well plate at desired timepoint (usually 5 days following radiation unless otherwise stated) for 15 minutes using 4% PFA (w/v, in PBS) followed by washing 3 times with PBS. Cells were then permeabilised using 0.2% Triton X-100 (v/v, PBS) for 10 min and then washed twice with PBS to halt permeabilisation. Non-specific antibody binding was blocked by incubation with a blocking solution for 1 hour at RT. Blocking solution contained 1% BSA (w/v, PBS) supplemented with 1X Fish Skin Gelatin (v/v, PBS). Primary antibodies (**Supplementary Table 3**) were diluted in blocking solution and cells incubated with primary antibody solution for 1 hour at RT. Following incubation, primary antibody was then removed by washing 3 times with PBS. Secondary antibodies (**Supplementary Table 4**) conjugated to Alexa-594 or Alexa-488 fluorophores were then diluted in blocking solution added to wells to be incubated in dark for 1 hour. Secondary antibody was then removed by washing 3 times with PBS and nuclei counterstaining with 1µg/mL DAPI (w/v, PBS) for 10 minutes. Cells were then washed with PBS three times. Immunofluorescence image acquisition was performed using an automated Opera Phenix high-throughput confocal microscope (Perkin Elmer).

Wells were imaged using a 10x, 20x or 40x water objective binning of images were used to reduce image file sizes. Fluorophores were imaged using pre-set ‘DAPI’, ‘AlexaFluor 594’ & ‘AlexaFluor 488’ wavelengths on microscope respectively. Depending on staining at least 20 fields per well were captured for 10x, 20x and 40x objectives respectively. High content image analysis was carried out using the Opera Phenix Harmony software (Perkin Elmer). DAPI nuclear counterstain was used to segment cells.

### LysoTracker staining

For live imaging, Lysosomes were visualised by incubation of 100 nM LysoTracker Deep Red (Molecular Probes, Thermo Fisher) for 30 minutes in complete media. Cells were counterstained with Hoescht for 10 minutes at a concentration of 1ug/ml, diluted in complete media. Fluorophores were imaged using pre-set ‘DAPI’, ‘AlexaFluor 647’ wavelengths on the microscope respectively. 10 fields per well were captured using a 40x water objective. High content image analysis was carried out using the Opera Phenix Harmony software (Perkin Elmer). Hoescht nuclear counterstain was used to segment cells. Cytoplasmic fluorescence intensity was measured per nuclei.

### Meso Scale Discovery (MSD) for cytokine and chemokine detection in cell culture supernatant

Supernatant was centrifuged before use and stored at −80°C. Cytokine concentrations (pg/ mL) were normalized to cell counts in each well (cell number/mL) resulting in cytokine concentration/cell (pg/cell). Data was log transformed for visual representation. Cytokine levels were measured using a Meso Scale Discovery multiplex kit (Meso Scale Diagnostics, Rockville, Md), according to the manufacturer’s instructions. IFN-γ, IL-1β, IL-2, IL-4, IL-6, IL-8, IL-10, IL-12p70, IL-13 and TNF-α were measured by V-PLEX MSD assay, and MIG (CXCL9) and IL-18 were measured by U-PLEX MSD assay. The plates were analyzed on the MSD instrument (QuickPlex SQ120). Concentrations were log transformed for visual representation.

### Transduction of primary human DMG cell lines

The plasmid pBM N(CMV-copGFP-Luc2-Puro) was a gift from O.Williams (Addgene plasmid 80389, RRID:Addgene_80389); and pMD2.G and psPAX2 were gifts (Addgene plasmid 12259, RRID:Addgene_12259; and Addgene plasmid 12260, RRID:Addgene_12260). HSJD-DIPG007 and SU-DIPG-VI cells were used for transduction. Cells were seeded in 24 well plates, and transduced at 70% confluence. Medium was removed from the tissue culture plate by aspiration and replaced with fresh complete medium containing 5-8 µg/ml polybrene (Millipore). Viral supernatant was added to achieve a multiplicity of infection (MOI) of 25%. After 18 hours, the medium was replaced with fresh complete medium to remove lentivirus and polybrene. Transduced cells were expanded and sorted using flow cytometry assisted cell sorting (FACS), based on GFP fluorescence sorted cells were expanded and stored in liquid nitrogen.

### Dose response cell assays

Cells were plated at a density of 5,000-10,000 cells/well on laminin-coated 96-well plates (black opaque) in a minimum of triplicates. Cells were first irradiated using IBL 437C (CIS Bio-International, Codolet, France) at doses stated (6 Gy, 12 Gy, 24 Gy, 36 Gy). Cells were allowed to sit for 18 hours, following this media was removed completely, and compounds were added to each well and incubated at 37°C in 5% CO2, 95% humidity for 3 days (72 hours. Cell viability was assessed by the CellTiter-Glo luminescent cell viability assay (Promega). SynergyFinder (https://synergyfinder.fimm.fi) was used for interactive analysis and visualization of drug combination profiling data following the Bliss independence model^32^. To calculate drug sensitivity scores a publicly available online platform was used, Breeze. Breeze uses a dose–response curve fitting process to directly translate raw cell viability measurements to clinically interpretable dose values. The fitted dose–response curve is then used to summarise and quantify the observed response into a single metric, such as drug sensitivity score (DSS)^33^.

### Assessment of apoptosis

Caspase 3/7 activity assay was determined using 100µl Caspase-Glo® 3/7 Reagent (Cat.# G8090), which was added directly to the cells in 96-well plates and incubated for 1 hour before recording luminescence on a Microplate luminometer - GloMax® Navigator - Promega. Each point represents the average of 3 wells. The “no cell” blank control value has been subtracted from each. Annexin V cell viability analysis used the Annexin V Fluorescent Reagent (Essen Bioscience), which was added to the media, together with vehicle (0.1% DMSO) or 0.1 µM Navitoclax for 48 hours. Images were collected every 2 h with an Incucyte® S3 Live-Cell Analysis System microscope (Essen Bioscience) over time. A total of 5 pictures per well were analysed using the IncuCyte ZOOM™ software analyser, and total and annexin V-positive cells were counted using ImageJ software.

### Generation of PDX models

A single-cell suspension (eGFP-LUCF-SU-DIPG-VI) was prepared immediately before implantation in 8-10 weeks old NOD-SCID male mice (Charles River). For intracranial procedures, animals were anesthetised with isoflurane. The cranium was exposed via midline incision under aseptic conditions and 1×1mm deep hole was made through the skull to the dura. Mice were placed in a stereotactic apparatus and 250,000 cells in 2-3 µL were stereotactically implanted in the pontine area using a 26SG Hamilton syringe using a rate of ∼ 1 µL/min. Co-ordinates used were 1.0mm lateral to midline, 0.8mm posterior to lambda, and –4.2 mm deep to cranial surface. A small pocket was created for the cells, which were injected −3.2 mm deep to the cranial surface. At the completion of infusion, the syringe needle was allowed to remain in place for a minimum of 2 minutes, then slowly manually withdrawn to minimize backflow of the injected cell suspension. The skin was closed with VetBond™ Adhesive. Mice were weighed daily for one week, and dosed for 48 hours with Metacam, at 5 mg/kg.

### Mouse brain irradiation

An Xstrahl Small Animal Radiation Research Platform (SARRP, Xstrahl Inc., Suwanee, GA) S/N 525722 irradiator (225 kV X-ray tube, half value layer 0.847 mm Cu), with 0.1 mm integral Be filtration, was used for mouse treatments. Cone-beam CT was performed before each treatment to confirm target position. Each mouse was anaesthetised with isoflurane and positioned for optimal brain targeting on a 3D printed bed. The bed was rotated between the X-ray source and a digital flat-panel detector. The images were obtained at 60 kVp and 0.8 mA with 1 mm Al filtration of an uncollimated primary beam. During rotation, 360 projections were acquired (approximately 1° increments for each projection, approximately 0.01 Gy total radiation dose). The cone-beam CT projections were rendered into a 3D image reconstruction, using the FDK algorithm with a voxel size from between 0.01 and 5 mm. Muriplan software was used to set radiodensity thresholds for different tissues and enable treatment planning, setting an iso-centre to target the injection site of the brain individually. Dose calculation was computed using a Monte Carlo simulation superposition-convolution dose algorithm, similar to those used clinically.

Mice received a total of either 12 or 15 Gy in 2-3 Gy fractions, delivered as an arc, targeted to maximise delivery to the whole brain (excluding the olfactory bulbs). The beam used was 220 kV and 13 mA, filtered with 0.15 mm Cu, dose rate 2.37 Gy min−1 under reference conditions. The beam was collimated, to match the treatment volume, with beam cross section up to 10 × 10 mm2. The time calculated to deliver 2-3 Gy to individual mice varied between at a dose rate of 0.037 Gy/sec with an exposure time of 50s to a designated field. Tumour growth was monitored using an IVIS Spectrum in bivo imaging system (Perkin Elmer). A total of 10 minutes following s.c injection of D-luciferin (Perkin Elmer) bioluminescent images were acquired under isoflurane anaesthesia. Tumour size was quantified by calculating total flux (photons/sec) using IVIS software.

## Results

### H3K27M-altered DMG cancer cell lines undergo senescence induction when exposed to ionising radiation

We first sought to test whether H3K27M-altered DMG cell lines of different genotypes (**Supplementary Table 1**) could be induced to senescence by irradiation. Three cell lines (ICR B117, HSJD-DIPG007 and SU-DIPG-IV) were exposed to single dose of ionising irradiation ranging from 12, 24 and 36 Gy and analysed at different time points for EdU incorporation and senescence-associated beta-galactosidase (SA-β-Gal) staining, markers of cellular proliferation and senescence, respectively (**Figure 1A**). Surviving tumour cells showed reduced EdU incorporation and increased SA-β-Gal staining, consistent with the induction of senescence response, in a dose and time dependent manner, relative to non-irradiated controls (**Figure 1, B-E**). While a dose of 8 Gy or below did not cause significant changes to EdU incorporation or SA-β-Gal staining (data not shown). This arrest in the cell cycle following ionising radiation was found to be transient following a dose of 12 Gy, with cells re-entering the cell cycle after ∼21 days. However, doses of either 24 or 36 Gy were able to induce a stable proliferative arrest (**Figure 1F**). DMG patients are clinically treated with 54 Gy in 30 fractions, of 1.8 Gy^34^. To attempt to mirror this more clinically applicable fractionation schedule, we exposed H3K27M-altered DMG cancer cells to 24 Gy, in 12 fractions of 2 Gy (Supplementary Figure 1A). This regime led to a similar response to that observed after exposure of the cells to a single dose, i.e., an increase of SA-β-Gal staining and a decrease in EdU incorporation (**Supplementary Figure 1B, C**). Moreover, qRT-PCR analysis revealed that cell irradiation led to the upregulation of *CDKN1A* (encoding p21) expression and down-regulation of *LMNB1*, encoding Lamin B1, a component of the nuclear lamina that is downregulated in senescent cells in a variety of cell contexts^35,36^ (**Supplementary Figure 1D).** These data suggest that irradiation, either in single doses or in a more clinically relevant fractionated schedule, results in a stable growth arrest, consistent with the induction of cellular senescence

**Figure 1.**
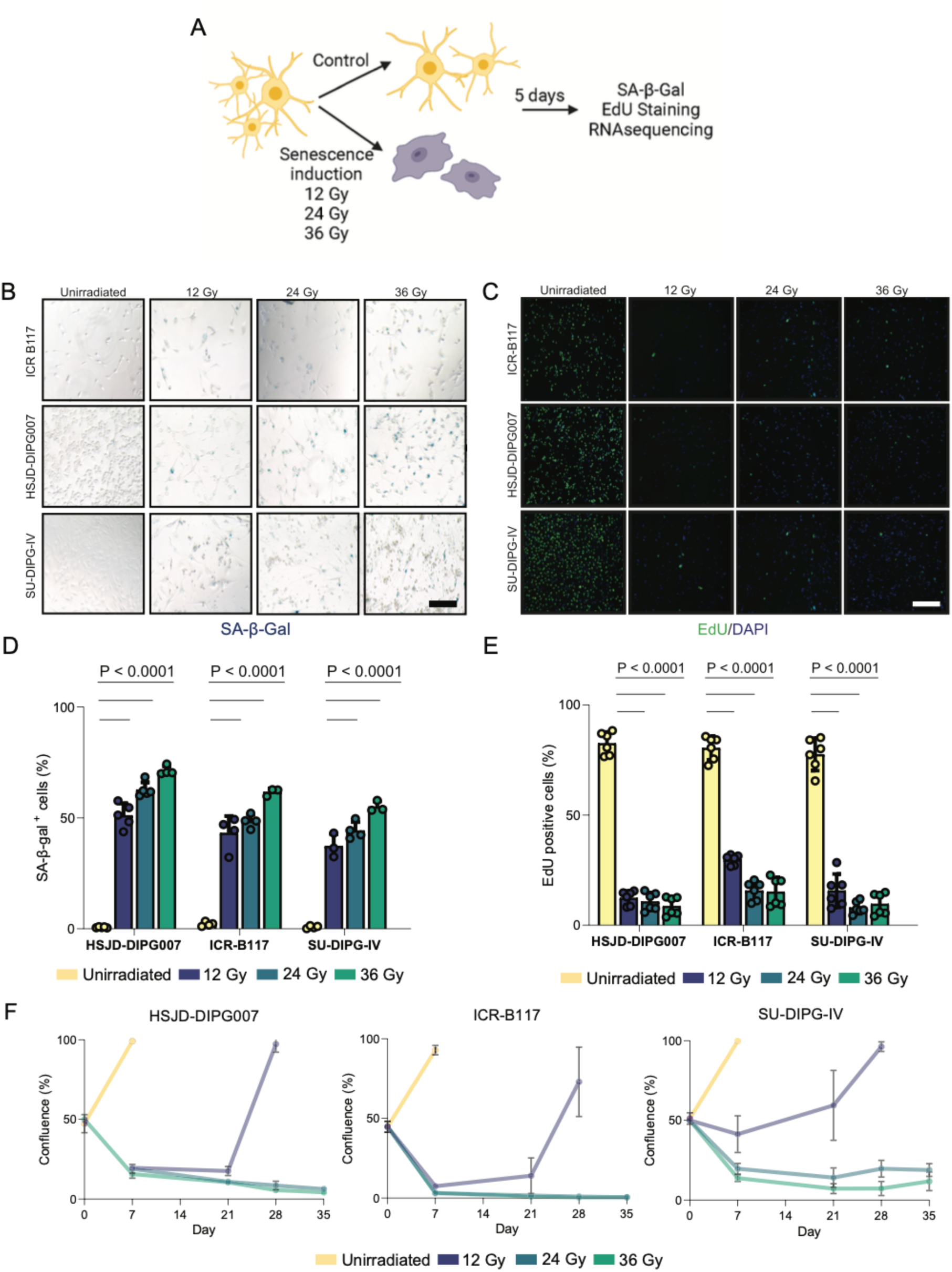
H3K27M-DMG cells following radiation exhibit growth arrest and express markers of senescence. (A) Schema outlining the experimental protocol for senescence induction with radiation. (B) Representative images of SA-β-gal staining of unirradiated control and irradiated DMG cells after different doses of radiation. (C) Representative images of EdU staining of unirradiated control and irradiated DMG cells after different doses of radiation. (D) Quantification of SA-β-gal positive cells from experiments shown in B. (E) Quantification of EdU positive cells from experiments shown in C. (F) Cell confluency of DMG cell lines either unirradiated or treated with single doses of radiation (12, 24, 36 Gy). Cells were monitored over a 35-day period. Error bar mean±SD. Colour relating to treatment. Legend colour representation of doses of radiation used. One-way ANOVA. Scale bar = 50µM.

To characterise further this potential senescent phenotype, we used a combination of biochemical and cytological staining techniques. Relative to unirradiated control cells, irradiated DMG cell lines displayed increased expression of the cyclin-dependent kinase inhibitors, p21, p27 (encoded by the *CDKN1B*) and p57 (encoded by the *CDKN1C* gene) following immunofluorescence analysis (**Figure 2A, B and Supplementary Figure 2A, B**). Conversely, phosphorylated Rb (pRB) protein expression, an important regulator of cell cycle progression, was downregulated in irradiated cells relative to non-irradiated control cells (**Figure 2A**). Irradiation is expected to cause DNA damage, which is a potent senescence inducer^37^. γH2AX immunofluorescence staining, which detects double DNA breaks^38^, revealed a significant increase in its expression in irradiated relative to non-irradiated control cells (**Figure 2A, B**). Moreover, CRYAB expression, a member of the small heat shock family of proteins and marker of cellular senescence^39^, was also upregulated in irradiated DMG cells (**Supplementary Figure 2C, D**). Another key feature of senescent cells is the enlargement of the lysosomal compartment^36^. Staining of the cells with Lysotracker, a red-fluorescent dye that labels lysosomes, revealed a significant increase in fluorescence intensity in irradiated relative to non-irradiated cells, indicating lysosomal expansion **(Figure 2A, C**). Senescence induction upon irradiation was also demonstrated in an additional DMG cell line, HSJD-DIPG14A (**Supplementary Figure 3**). Together, these analyses suggest that irradiation leads to a senescent phenotype in both TP53 mutant (ICR-B117, SU-DIPG-IV) and TP53 wild-type (HSJD-DIPG007, HSJD-DIPG14) H3K27M-altered DMG cell lines.

**Figure 2.**
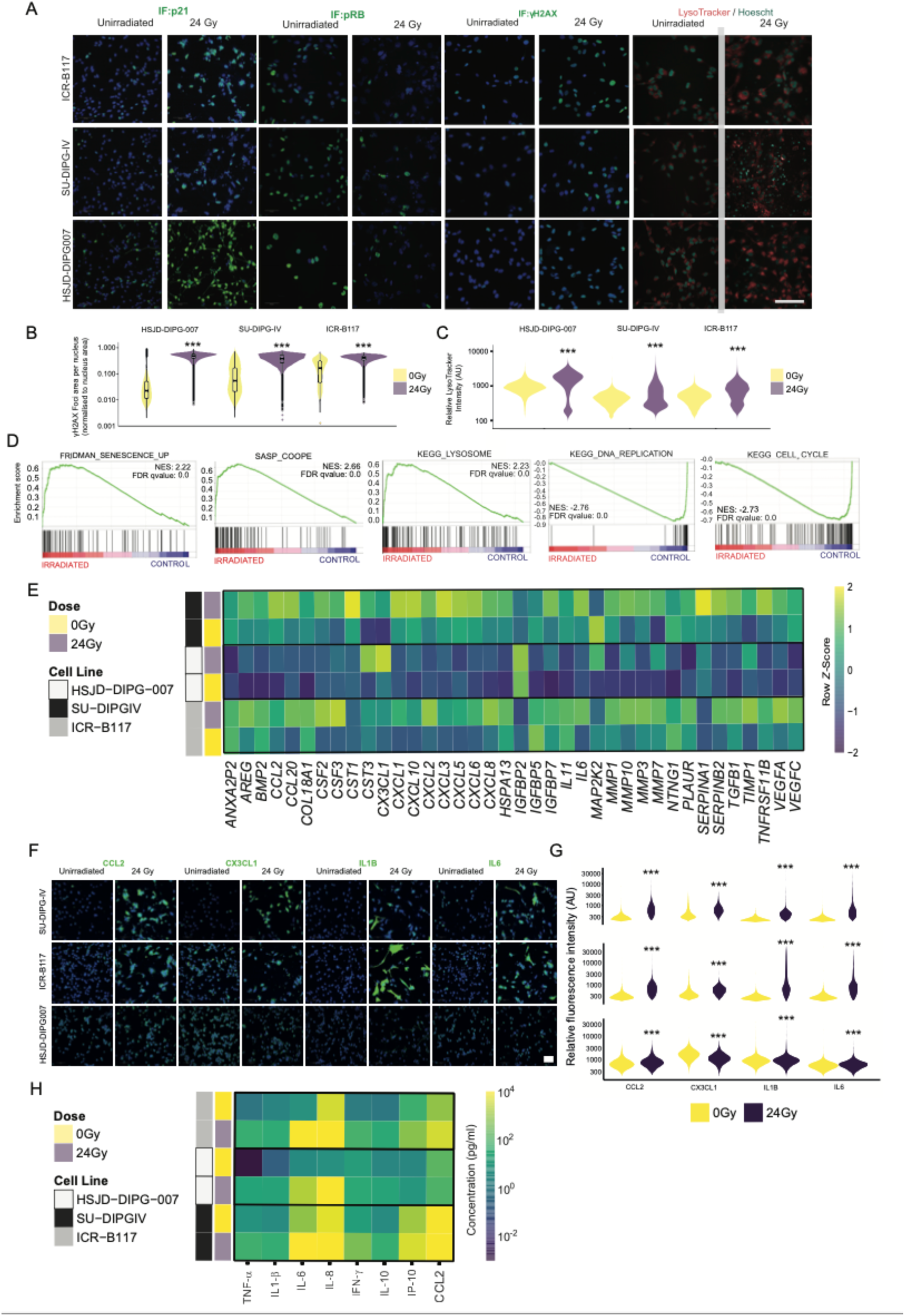
H3K27M-DMG cells following radiation exhibit a robust senescence phenotype. (A) Immunofluorescence (IF) staining in three DMG cell lines following single dose of irradiation (24 Gy) with p21 (CDKN1A), phospho-RB and γH2AX, and red-fluorescent dye LysoTracker live cell staining. Scale bar = 100µM (B, C) Quantification of γH2AX and LysoTracker stainings shown in A. Fluorescence intensity is indicated in arbitrary units (AU). (D) Gene set enrichment Analyses showing a significant enrichment for the FRIDMAN_SENESCENCE_UP, SASP_COOPE and KEGG_LYSOSOME gene sets in irradiated DMG versus unirradiated control cells. Note that the KEGG_DNA_REPLICATION and CELL_CYCLE gene sets are enriched in unirradiated control cells. (E) Heatmap of RNA sequencing analysis of DMG cell lines following 24 Gy of radiation revealing an upregulation of genes associated with senescence/SASP. (F) Immunofluorescence staining of DMG cell lines following irradiation (24 Gy) against the SASP factors: CCL2, CX3CL1, IL1B and IL6. (G) Quantification of fluorescence intensity (arbitrary units, AU) is plotted for each cell line. (H) Meso-scale discovery cytokine quantification was carried out on conditioned media 5 days following irradiation. Heatmap represents log fold change cytokine concentration (pg/ ml). One-way ANOVA. NES = normalised enrichment score. FDR = false discovery rate.

To complement our cytological analyses with a molecular characterisation, we performed bulk RNA-sequencing on ICR-B117, SU-DIPG-IV and HSJD-DIPG007 cell lines prior to and after irradiation (single dose of 24 Gy). *In silico* analysis combining the datasets from the three DMG cell lines revealed that a total of 2863 genes were significantly differentially expressed between the irradiated and non-irradiated cell lines (adjusted P value ≤0.01, Log2FC ≥±2), comprising 1452 upregulated genes and 1411 downregulated genes (**Supplementary Figure 2E, F and Supplementary Table 5**). Among the significantly upregulated genes in irradiated cells, we found a number of genes relevant to the senescent phenotype: (i) cell cycle inhibitors (e.g. *CDKN1A* (encoding p21, Log2FC 1.68); (ii) lysosomal genes (e.g. *GLB1, MAN2B, HEXA*; Log2FC 0.57, 1.21, 0.96, respectively); (iii) SASP factors (e.g. *MMP3* (Log2FC; 6.29), *IL1B* (Log2FC 8.28), *CXCL8* (Log2FC: 8.07) *CX3CL1* (Log2FC; 5.11) and *FGF7* (Log2FC; 6.43). In agreement with a senescence phenotype, genes normally associated with proliferation, such as *E2F2, MYBL2,* and *ZWINT* were downregulated in irradiated cells (Log2FC: −2.6, −2.65, −2.39, respectively), as well as *LMNB1* (Log2FC; −2.24) (**Supplementary Figure 2E-H**).

Gene set enrichment analysis (GSEA) revealed a significant enrichment for senescence^40^, SASP^41^ and KEGG Lysosome signatures in irradiated compared with non-irradiated control cells (**Figure 2D**). Conversely, gene expression signatures associated with cell cycle progression were all significantly enriched in non-irradiated control cells (**Figure 2D**). GSEA of the Hallmark gene set collection (from the Molecular Signatures Database (MSigDB)) revealed enrichment of stress and inflammation related processes (i.e., Hypoxia, IL6-JAK-STAT3 signalling, inflammatory response and TNFα signalling) in irradiated DMG cells (**Supplementary Figure 2I**) and proliferative responses (i.e., Mitotic spindle, G2M checkpoint and E2F targets) in the unirradiated control DMG cells (**Supplementary Figure 2J**). In agreement, gene ontology (GO) analysis revealed similar stress and inflammatory processes in irradiated cells (**Supplementary Figure 2K**) and proliferative responses in unirradiated cells (**Supplementary Figure 2L**).

From our transcriptomic analysis we also observed upregulation of a number of SASP factors in irradiated DMG cell lines (**Figure 2E**). To validate these findings, we performed immunofluorescence staining against CCL2, CX3CL1, IL1β and IL6, in either irradiated or unirradiated control DMG cells, revealing a significant upregulation of these factors in the irradiated group (**Figure 2F, G**). Moreover, ELISA analysis of conditioned medium demonstrated the increased secretion of several senescence-associated cytokines and chemokines in irradiated DMG cells relative to control medium, including the prominent SASP factors IL1β, TNFα, IL6 and CCL2 (Coppé, Patil et al. 2008) (**Figure 2H**). Importantly, clinically relevant fractionated doses of ionizing radiation were also sufficient to induce upregulation of the SASP markers IL1B and CCL2, relative to non-irradiated proliferative control cells (**Supplementary Figure 1D**).

Collectively, these molecular and cellular analyses demonstrate that ionising irradiation leads to DNA damage, decreased proliferation, induction of senescence markers and activation of a secretory phenotype in H3K27M-altered human DMG cell lines independently of the mutational status or *TP53*.

### Navitoclax is a potent senolytic in H3K27M-altered human DMG cell lines

Having revealed a senescent phenotype in response to radiation, we next aimed to explore possible therapeutic implications of leveraging the phenotype through the use of seno-therapies to ablate senescent DMG cells. We tested four described senolytic agents, Dasatinib plus Quercetin (D+Q), Piperlongumine, Alvespymicin and Navitoclax, on senescent (irradiated) and control proliferative (unirradiated) DMG cells.

Dose response curves for D+Q, Piperlongumine and Alvespymicin did not reveal the selective killing of senescent relative to proliferative DMG cells in any of the four cell lines tested (HSJD-DIPG14A, HSJD-DIPG007, SU-DIPG-IV and ICR-B117) (**Supplementary Figure 4**). In contrast, Navitoclax showed a potent and selective efficacy on HSJD-DIPG14A, HSJD-DIPG007 and SU-DIPG-IV senescent cells with IC50s of 0.01 µM, 0.04 µM, and 0.07 µM, respectively (**Figure 3A**; **Table 1**). The cell line ICR-B117 showed resistance to Navitoclax, relative to the other DMG cell lines, and the IC50 was 0.34 µM.

**Figure 3.**
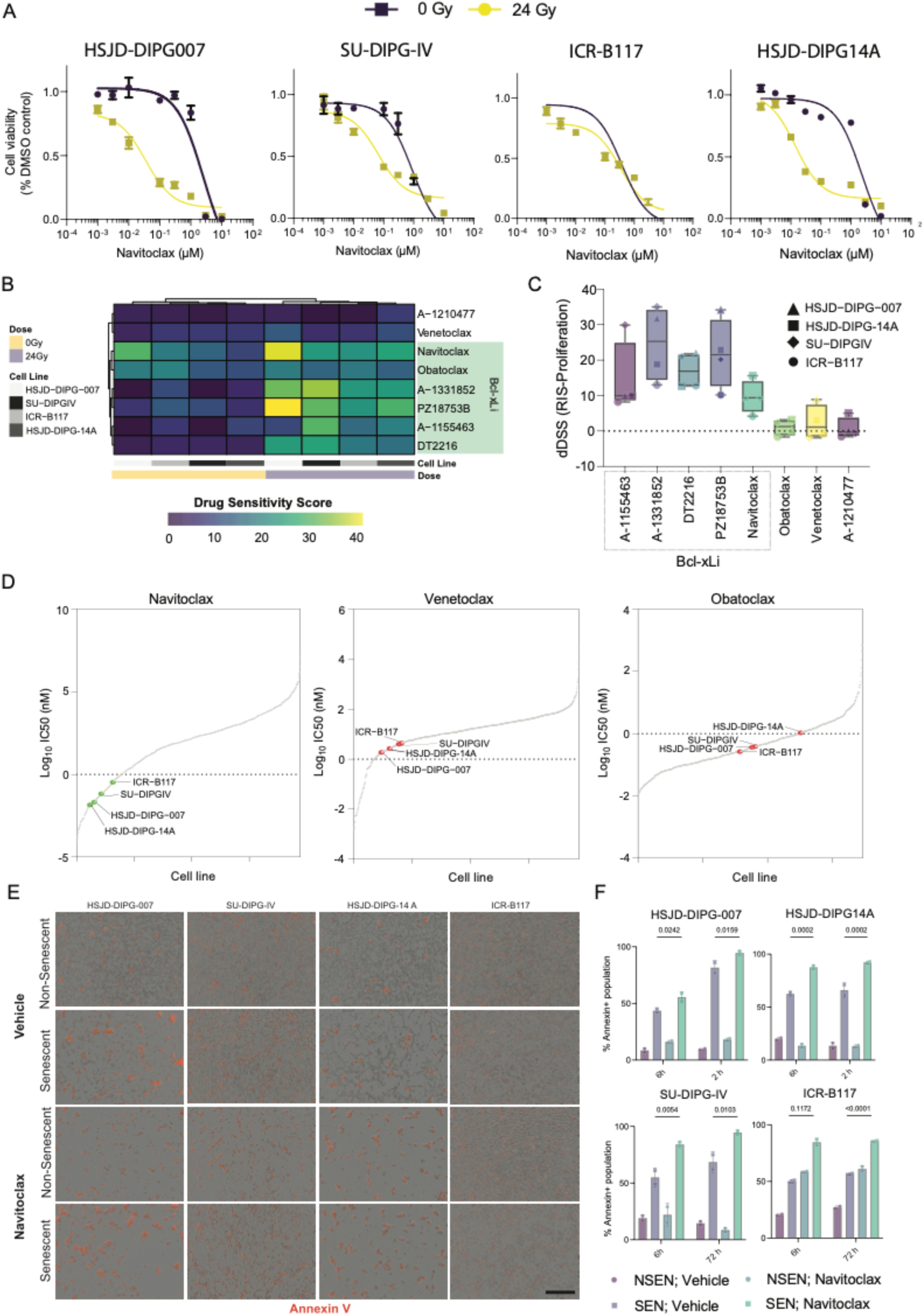
Bcl-xL inhibition is a targetable vulnerability in H3K27M-DMG cells following radiation. (A) Dose-response curves of four different human DMG cell lines exposed to a range of Navitoclax concentrations in senescent (yellow) or proliferating (purple) states. Values are shown are mean ± SD of three technical replicates. (B) Drug sensitivity scores are calculated based on dose-response data (n=3 biological replicates for each drug, cell line and state) using Breeze software. (C) Dotplot depicting mean differential drug sensitivity scores (dDSS) of the indicated drugs across four DMG cell lines (mean±SD). dDDS are calculated by subtracting DSS of proliferating cells from DSS of senescent cells. (D) Ranking of IC50 values of BH3 mimetics sensitivity of the four senescent DMG cell lines against 967 cell lines from the GDSC database. The sensitivity line is set at 1µM. (E) Representative images of Annexin V staining in DMG cells in the proliferating (unirradiated) and senescent (24 Gy irradiation) states exposed to navitoclax (10 µM) or vehicle (0.1% DMSO) for 6 hours post treatment. Scale bar at lower magnification = 400 µm. (F) Quantification of Anexin V-expressing proliferating and senescent DMG cells at 6 and 12 hours of Navitoclax or vehicle exposure. Data represent mean ± SD. Statistical significance was calculated using a two-tailed student’s t test.

**Table 1.** IC50 values from proliferative and senescent H3K27M-altered human DMG cell lines cultured in the presence of anti-apoptotic protein inhibitors.

|  |  | IC50 (μM) |  |  |
| --- | --- | --- | --- | --- |
| Drug | Cell lines | Proliferative cells | Senescent cells | Senolytic index* |
| Navitoclax | HSJD-DIPG007 | 2.55 | 0.04 | 63.7 |
| Venetoclax |  | 26.26 | 1.87 | 11.9 |
| Obatoclax |  | 0.30 | 0.36 | 0.5 |
| PZ18753B |  | 7.81 | 0.02 | 390.5 |
| DT2216 |  | 26.19 | 0.10 | 261.9 |
| Nav-gal |  | 3.74 | 0.10 | 37.4 |
| Navitoclax | HSJD-DIPG14A | 2.37 | 0.01 | 237 |
| Venetoclax |  | 3.62 | 2.66 | 1.3 |
| Obatoclax |  | 1.89 | 1.07 | 1.7 |
| PZ18753B |  | 18.18 | 0.02 | 909 |
| DT2216 |  | 0.56 | 0.09 | 6.2 |
| Nav-gal |  | 1.31 | 0.15 | 8.7 |
| Navitoclax | SU-DIPG-IV | 0.90 | 0.07 | 12.8 |
| Venetoclax |  | 3.85 | 4.00 | 0.9 |
| Obatoclax |  | 1.04 | 0.39 | 2.6 |
| PZ18753B |  | 4.81 | 0.01 | 481 |
| DT2216 |  | 0.33 | 0.01 | 33 |
| Nav-gal |  | 39.62 | 0.15 | 246.1 |
| Navitoclax | ICR-B117 | 0.38 | 0.34 | 1.1 |
| Venetoclax |  | 3.18 | 4.40 | 0.7 |
| Obatoclax |  | 0.25 | 0.27 | 0.9 |
| PZ18753B |  | 0.63 | 0.14 | 4.5 |
| DT2216 |  | 0.87 | 0.06 | 14.5 |
| Nav-gal |  | 1.66 | 0.14 | 11.8 |

\* Senolytic index: IC50 proliferative cells/IC50 senescent cells

Navitoclax can inhibit Bcl-2, Bcl-xL, and Bcl-w^42^. In order to elucidate the specific protein responsible for the senolytic effect of Navitoclax, we used several BH3 mimetics, including both chemical probes and clinically approved compounds, with specific inhibitory profiles (**Supplementary Table 2)**. Drug sensitivity scores (DSS) were determined from dose-response curves data obtained in irradiated and non-irradiated, proliferative control cells (**Supplementary Figure 5**). Our findings revealed low DSS values (DSS Range: 5.8-12.5) for Venetoclax, a Bcl-2 inhibitor, and A-1210477, an MCL-1 inhibitor. These low DSS values indicate low susceptibility for these inhibitors in our experimental setup (**Figure 3B**). On the other hand, DSS values obtained for all inhibitors with strong affinity to Bcl-xL (Navitoclax, A-1331852, A-1155463, PZ18753B, DT2216) indicated higher sensitivity for all cell lines in the senescent state (DSS Range: 9.9 - 41) (**Figure 3B**). Furthermore, the differential DSS (dDSS), obtained from subtracting the DSS of non-irradiated (proliferative cells) from the DSS of radiation-induced senescent cells (RIS-DSS) were positive for all cell lines treated with Bcl-xL inhibitors, indicating that senescent DMG cells exhibited a preferential susceptibility to Bcl-xL inhibition rather than to Bcl-2 or MCL-1 inhibition (**Figure 3C**).

Next, we compared the IC50 values obtained from the Genomics of Drug Sensitivity in Cancer (GDSC) study^43^, which encompasses 967 cell lines treated with drug compound libraries, with the IC50s observed in DMG cell lines for three different BH3 mimetics, Navitoclax, Venetoclax and Obatoclax (**Figure 3D**). The IC50s of Navitoclax in irradiated DMG cells were lower than 0.34 µM, ranking among the top 16% of most sensitive cell lines. Specifically, HSJD-DIPG-14A, HSJD-DIPG007, SU-DIPG-IV and ICR-B117 ranked 58^th^, 76^th^, 107^th^ and 157^th^ out of the 967 cell lines of reference (**Figure 3D**). In contrast, the IC50s of Venetoclax were higher in senescent DMG cells (1.87-4.4 µM), and these 4 DMG cell lines ranked 119^th^ (HSJD-DIPG007), 154^th^ (HSJD-DIPG-14A), 194^th^ (SU-DIPG-IV) and 205^th^ (ICR-B117), clearly demonstrating a lower sensitivity compared with Navitoclax (**Figure 3D**, **Table 1**). Senescent DMG cells were also less sensitive to obatoclax (ICR-B117: 394^th^; HSJD-DIPG-007: 441^st^; SU-DIPG-IV: 453^th^; HSJD-DIPG-14A: 628^th^) with IC50 of 0.27-1.07 µM (**Figure 3D**, **Table 1**).

Finally, we sought to ascertain whether Navitoclax ablates the senescent DMG cells by apoptosis. Senescent (irradiated) and proliferative (non-irradiated) DMG cells were treated with 0.1µM of Navitoclax or vehicle and caspase 3/7 activity quantified using either a luminescent assay or an automated real time measurement of Annexin V staining in a live-cell analysis system. Treatment with Navitoclax resulted in increased Annexin V signal and caspase activity, suggesting induction of apoptosis (**Figure 3E, F; Supplementary Figure 6**. Together, these analyses demonstrate that senescent H3K27M-altered human DMG cells are particularly sensitive to apoptotic induction through Bcl-xL inhibition.

### Proteolysis-Targeting Chimera (PROTAC)-mediated Bcl-xL degradation and galacto-conjugated Navitoclax are senolytics in senescent H3K27M-altered human DMG cells

A potential limitation of using navitoclax clinically is the on-target toxicity affecting platelets, which leads to significant thrombocytopenia ^44,45^. However, there have been two novel strategies to circumvent possible toxicity through the use of Proteolysis-Targeting Chimera (PROTAC)-mediated Bcl-xL degradation^46^ and a galacto-conjugated form of Navitoclax that is specifically activated in senescent cells (Nav-Gal)^22^. Therefore, we sought to test the efficacy of Nav-Gal and two different PROTACs, named DT2216, targeting only Bcl-xL^46^ and PZ18753B^47^, targeting both Bcl-xL and Bcl-2, which have recently been shown to be effective senolytics *in vitro* and *in vivo*.

To assess the senolytic capability of these compounds, dose response data were used to calculate the IC50s on senescent and proliferative cells (**Figure 4**; **Table 1**). To compare these IC50s in a meaningful way, we decided to calculate the senolytic index (SI), defined as the ratio between the IC50 values of proliferative and senescent cells. The SIs ranged from 8.54-264.66 μM (Nav-Gal), 45.93-266.16 μM (DT2216), 4.48-1163.89 μM (PZ18753B) and 1.13-165.78 μM (Navitoclax), suggesting that Nav-Gal and PROTACs exhibit a greater selectivity for targeting senescent over proliferative cells.

**Figure 4.**
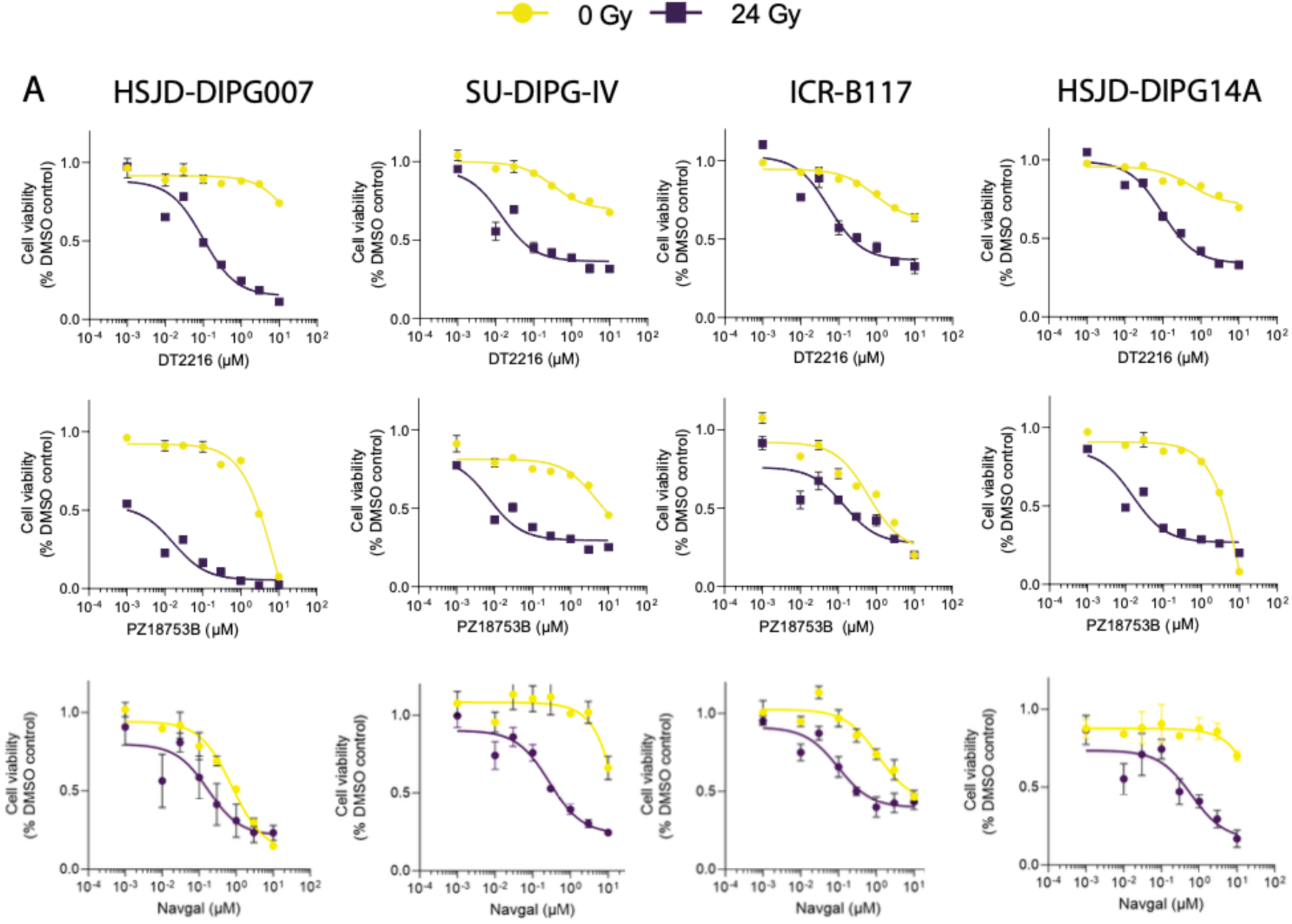
The prodrug Nav-Gal and Bcl-xL targeting PROTACs shows efficient senolytic activity on senescent DMG cells. (A) Dose-response curves for the PROTACs DT2216 (targeting Bcl-xL) and PZ18753B (targeting Bcl-xL and Bcl-2), and the prodrug Nav-Gal on proliferating (yellow) and irradiated, senescent (purple) human DMG cells. Data is shown as mean ± SD.

### Strong synergistic effect of a combination therapy of irradiation with Navitoclax

As the standard of care for treating DMG patients is radiotherapy (RT), we aimed to assess the potential synergistic effect of combining radiation, which targets proliferative cells to induce cell death and senescence, with BH3-mimetics, which can ablate senescent cells. Cell viability was measured using Cell-Titre Glo in HSJD-DIPG-14A, HSJD-DIPG-007, SU-DIPG-IV and ICR-B117, 72 hours after combination of BH3-mimetic (Navitoclax, Obatoclax and Venetoclax) and increasing doses of radiation (0, 6, 12, 24, 36 Gy) **(Supplementary Figure 7A).** Combinatorial effects were calculated using the Bliss independence model since the treatments combined were assumed to act independently. In this model, scores ranging from −10 to +10 indicate an additive effect, whilst scores higher than +10 demonstrate synergy^48^. Overall, the most synergistic Bliss scores were obtained with a combination of RT with Navitoclax: HSJD-DIPG-14A, 22.1; HSJD-DIPG-007, 7.5; SU-DIPG-IV, 8.9; ICR-B117, 11.4 **(Supplementary Figure 7B).** The combination of RT with Obatoclax was the least effective (HSJD-DIPG-14A, 6.3; HSJD-DIPG-007, 1.1; SU-DIPG-IV, 1.8; ICR-B117, 2.1) **(Supplementary Figure 7B**). RT and Venetoclax combination led to a mixed response at inhibiting cell viability, HSJD-DIPG-14A, 18.3; HSJD-DIPG-007, 3.4; SU-DIPG-IV, 2.4; ICR-B117, 8.1 **(Supplementary Figure 7B).** Together with the data shown previously, these results confirm that H3K27M-altered human DMG cells show a strong sensitivity to a combination therapy of RT and Bcl-xL inhibition.

### Combination therapy of radiotherapy and Navitoclax reduces tumour burden and increases survival in a DMG PDX model

To assess *in vivo* the efficacy of Navitoclax and RT against DMG, we sought to generate xenograft models using the H3K27M-altered human DMG cells previously characterised. HSJD-DIPG-14A, SU-DIPG-IV and ICR-B117 cells have previously shown very low engraftment potential, and although HSJD-DIPG-007 cells engrafted with high efficiency in our hands, tumours showed little response to RT (data not shown). The cell line SU-DIPG-VI has previously been shown to develop DMG tumours in the pontine area upon transplantation^49^. We first confirmed that SU-DIPG-VI cells also show a senescent phenotype in response to irradiation and marked sensitivity to Navitoclax in the senescent state (IC50, 0.27 μM; Senolytic index, 6.3) (**Supplementary Figure 8).**

To enable the identification of tumours in vivo, we generated eGFP-LUCF-SU-DIPG-VI cells, which express eGFP and luciferase, and used these cells in orthotopic transplantations in NSG mice. A schematic of the preclinical experimental design is shown in **Figure 5A**. After cell transplantation into the pons, BLI was used as an estimate of tumour burden, and tumour-bearing mice distributed into 4 arms 20 days after transplantation (**Supplementary Figure 9**); (a) Untreated control group, which was not irradiated and was administered with the vehicle used to prepare Navitoclax; (b) Navitoclax-only treated group, was not irradiated and was administered Navitoclax (50 mg/kg/day), given via oral gavage 5 days a week for 4 weeks; (c) Radiotherapy group, which received 6 fractions of 2 Gy/day (12 Gy total (higher doses led to unacceptable toxicity)) and received the vehicle used to prepare Navitoclax; and (d) Combination therapy group, which was treated with both RT and Navitoclax as previously described (b and c).

**Figure 5.**
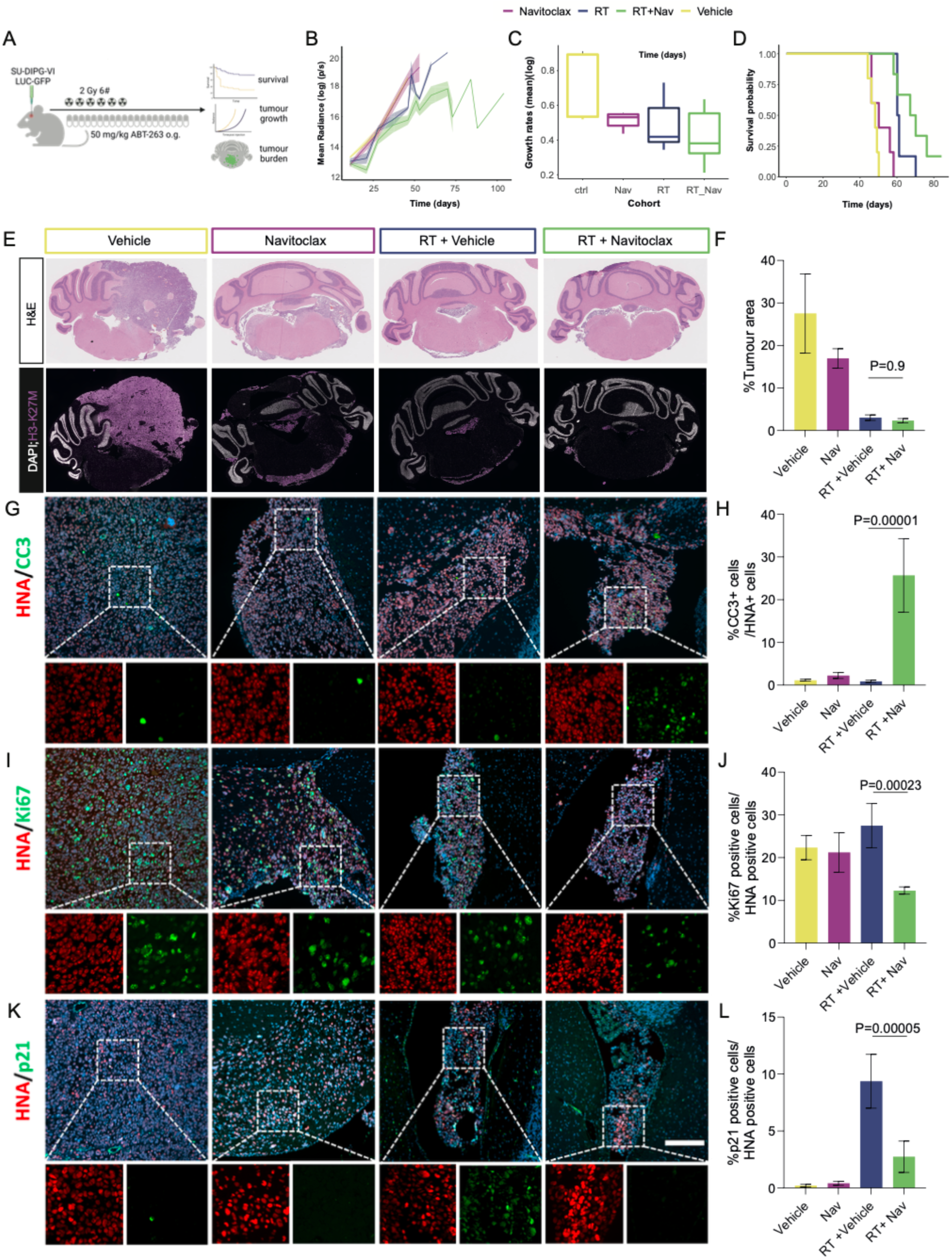
A combination therapy of Navitoclax and radiotherapy significantly decreases tumour burden and prolongs survival in a DMG orthotopic mouse model. (A) Schema outlining the *in vivo* experimental protocol. Upon orthotopic transplantation in NSG, tumour-harbouring mice were divided into four experimental groups (vehicle, Navitoclax, radiotherapy (RT)-treated and RT plus Navitoclax). (B) Growth rate of tumours using relative luciferase radiance is expressed as Δlog radiance/time. (C) Kaplan-Meier survival curves of mice in the fours experimental groups. (D) Representative histological images of brain sections of each experimental group stained for H3-K27M. (E) Quantification of the proportion of the brain section area occupied by H3-K27M-positive tumour cells from (D). (F, H and J) Double immunofluorescence staining of SU-DIPG-VI tumour bearing brain sections at end point for human nuclear antigen (HNA)-positive tumour cells with either p21Cip1 (P21) (F), Ki67 (H) or cleaved caspase 3 (CC3) (J). Scale bars; 100 µm. (G, I and K) Quantification of the proportion of HNA-expressing tumour cells which express the markers from (F, H and J, respectively). Data are mean±s.e. (stat = One Way ANOVA, P-value shown, n=3)

Analyses of the tumour growth rates (TGR, defined as BLI change over time) showed no significant differences between the navitoclax-only treated and untreated control vehicle groups (groups a and b) (**Figure 5B**). RT alone (group c) reduced TGR significantly in comparison with both, untreated vehicle control (group a, p=0.03) and Navitoclax only group (group b, p=0.002) (**Figure 5B**). Likewise, the combination of RT plus Navitoclax resulted in a significant reduction of TGR in comparison with groups a and b (p=0.02 and p=0.0008, respectively), and led to a trend towards a reduction that did not reach significance against the RT-alone group c (**Figure 5B**). Kaplan-Meier survival curves were generated to analyse the survival probability of the four groups of mice (**Figure 5C**). As with TGRs, significant differences in median survival were observed only between the RT-only (group c, 60.5 days) and the RT plus Navitoclax (group d, 68.5 days) compared with the vehicle (group a, 48 days) and Navitoclax-only (group b, 50 days) (RT-only: p=0.001 and p=0.001; RT plus Navitoclax: p= 0.001 and p=0.002). Although, the median survival was increased in the RT plus Navitoclax group (68.5 days) relative to the RT-only group (60.5 days), these differences did not reach significance (p=0.18).

Next, we assessed histologically and by immunofluorescence staining the tumours in the four experimental groups 10 days post-irradiation (**Figure 5D-K**). Quantification of the tumour area showed no significant differences between untreated vehicle control and Navitoclax-only treated groups (groups a and b) (**Figure 5D, E**). In contrast, treatment with RT alone or RT plus Navitoclax resulted in significant differences in tumour area relative to groups a and b (RT-only: p=0.0002 and p=0.03; RT plus Navitoclax: p=0.0001 and p=0.02). No significant differences were observed between the RT and the RT plus Navitoclax groups (**Figure 5D, E**). To assess whether RT was able to induce senescence, we performed double immunostaining against CDKN1A (p21), to detect senescent cells, and the human nuclear antigen (HNA), to reveal tumour cells (**Figure 5F**). These analyses showed a significant increase in the proportion of p21/HNA-double positive tumour cells in the RT-only group relative to the vehicle control (group a, p=1×10^−6^), Navitoclax-only group (group b; p=1×10^−6^), and importantly, the RT plus Navitoclax combination group (group d, p=5×10^−5^) (**Figure 5F, G**). These results are compatible with the notion, as previously shown in vitro in this study, that RT can induce a senescent response in DMG cells. Moreover, the significant reduction in p21-positive DMG tumour cells observed in vivo is compatible with the hypothesis that Navitoclax may cause the ablation of these RT-induced senescent tumour cells. To explore this hypothesis further, we performed double immunostaining against HNA and either cleaved-caspase-3, as a marker of apoptosis, or KI67, to detect cell proliferation (**Figure 5H, J**). The proportions of Ki67/HNA-double positive cells were not significant difference between groups a, b and c (vehicle control, Navitoclax-only control and RT-only control), however, the group treated with the combination therapy of RT plus Navitoclax showed a significant reduction in the Ki67 proliferative index (p=0.0002) (**Figure 5I**). Importantly, while the proportion of CC3/HNA-double positive DMG tumour cells in the vehicle control, Navitoclax-only treated and RT-only treated groups were almost negligible (around 0.9-2.3 %), there was a significant increase in numbers of double positive tumour cells in the RT plus Navitoclax group (25.7%; p=0.00001). Together, these analyses suggest that treatment with RT induces the appearance of p21-positive tumour cells which are sensitive to Navitoclax-mediated ablation.

## Discussion

In this study, we have demonstrated that ionising radiation results in both senescence induction (**Figures 1 and 2**) and increased sensitivity to BH-mimetics (**Figure 3**) in a set of H3K27M-altered human DMG cells. Using specific inhibitors, we show that senescent DMG cells’ survival depends mostly on the anti-apoptotic protein Bcl-xL rather than the family members Bcl-2 or Mcl-1 (**Figures 3B-D and 4**). We reveal that a combination of irradiation with Bcl-xL inhibitors result in the cell death of DMG cells in the senescent state (**Figure 3E, F**). However, not all the DMG cell lines tested exhibit the same sensitivity to Bcl-xL inhibition, and in particular ICR-B117 cells are more resistance (IC50 0.34 μM), while HSJD-DIPG-14A, HSJD-DIPG-007 and SU-DIPG-IV are more sensitive with IC50 values of less than 70 nM (**Table 1**). Senescence induction and sensitivity to Bcl-xL inhibition is supported by the data obtained in a PDX model of DMG (**Figure 5**). A fractionation regimen of 2 Gy per day for 6 days leads to significant tumour reduction, increased mouse survival and long-term accumulation of p21+ tumour cells. Moreover, combination of irradiation with Navitoclax results in significant anti-tumour response and reduction of both p21+ and Ki67+ tumour cells. Although our in vitro studies demonstrate a marked synergy between irradiation and Navitoclax, the in vivo paradigm used did not detect significant differences between the group treated with only irradiation and the irradiation plus Navitoclax groups. Nonetheless, there is a trend towards reduced tumour growth rate and increased mouse survival in the combination group compared with the irradiation-only group (**Figure 5B-D**).

Previous studies have shown that DMG cells can be induced into senescence by inhibition of BMI1 and EZH2^50–52^. Moreover, senescent cells induced through BMI1 inhibition are vulnerable to BH3-mimetics both in vivo and in vitro^51^, indicating the senescent response could be exploited therapeutically. Together with our data these findings hold high translational interest as compounds inhibiting Bcl-xL, such as Navitoclax, PROTAC, and Nav-Gal, are already in advanced clinical development, making them potential candidates for clinical phase I/II trials in paediatric DMG.

The sensitivity of senescent cells to Bcl-xL inhibition has also been shown in other brain tumours. The high-grade adult glioma, glioblastoma multiforme (GBM) is known for its aggressive and invasive nature, making it one of the most challenging brain tumours to treat. GBM can undergo senescence following radiation or temozolomide treatment and preclinical studies have shown that targeting senescent cells through Bcl-xL inhibition could be a valuable therapeutic strategy for improving GBM treatment outcomes^2,5,25,53^. Likewise, sensitivity to Bcl-xL inhibition has been observed in cell lines of pylocitic astrocytoma, the most common brain tumour in children, and adamantinomatous craniopharyngioma, the most common childhood pituitary tumour, suggesting a potential therapy against these tumours^28,30,54^. Recently, Venetoclax, also known as ABT-199, a Bcl-2 inhibitor, has been shown to cooperate with irradiation in controlling paediatric DMG^55^. Notably, in our study, the role of Venetoclax has not been found to be critical in maintaining survival of senescent, irradiated cells. Instead, we show that Bcl-xL inhibition is a more crucial target for senolytic activity. Nonetheless, treatment of senescent DMG cells with the dual PROTAC PZ18753B, which targets for degradation both anti-apoptotic proteins Bcl-xL and Bcl-2, results in the lowest IC50 values and highest senolytic scores in our study (**Table 1**), suggesting functional redundancy.

In summary, we show the efficacy of Bcl-xL inhibitors as senolytic compounds in senescent DMG cell lines, both in vivo and in vitro. The ongoing clinical development of less toxic and more selective Bcl-xL inhibitors and their evaluation in clinical trials for other tumours present exciting prospects for future therapeutic interventions in brain tumours, including DMG.

## Authorship

AV carried out most of the experiments. SH carried out the experiments shown in Figure 5D-K, and prepared all the figure panels for publication. RG, DG, JG, JA, JD contributed experimentally. DC, DZ, GZ, RM-M, AMC, CJ provided reagents. RC, JB, ML performed tumour imaging/irradiation. LD provided expertise in brain cell injection. DM, CJ and DH provided expertise and intellectual input. The project was designed by DH and JPM-B. JPMB wrote the paper, with help from SH and AV. All authors reviewed and approved this submission.

## Data availability

RNA sequencing data is available in accession number ()

## Acknowledgements

We thank core facilities such as the Pathology Department at the Institute of Neurology, the UCL Centre for Advance Imaging (CABI), UCL Genomics, UCL Biological Services Unit and CRUK City of London core facilities for facilitating this research. We thank our funders and sponsors for enabling the research presented in this manuscript.

## Supplementary Figures

**Suppl. Figure 1:**
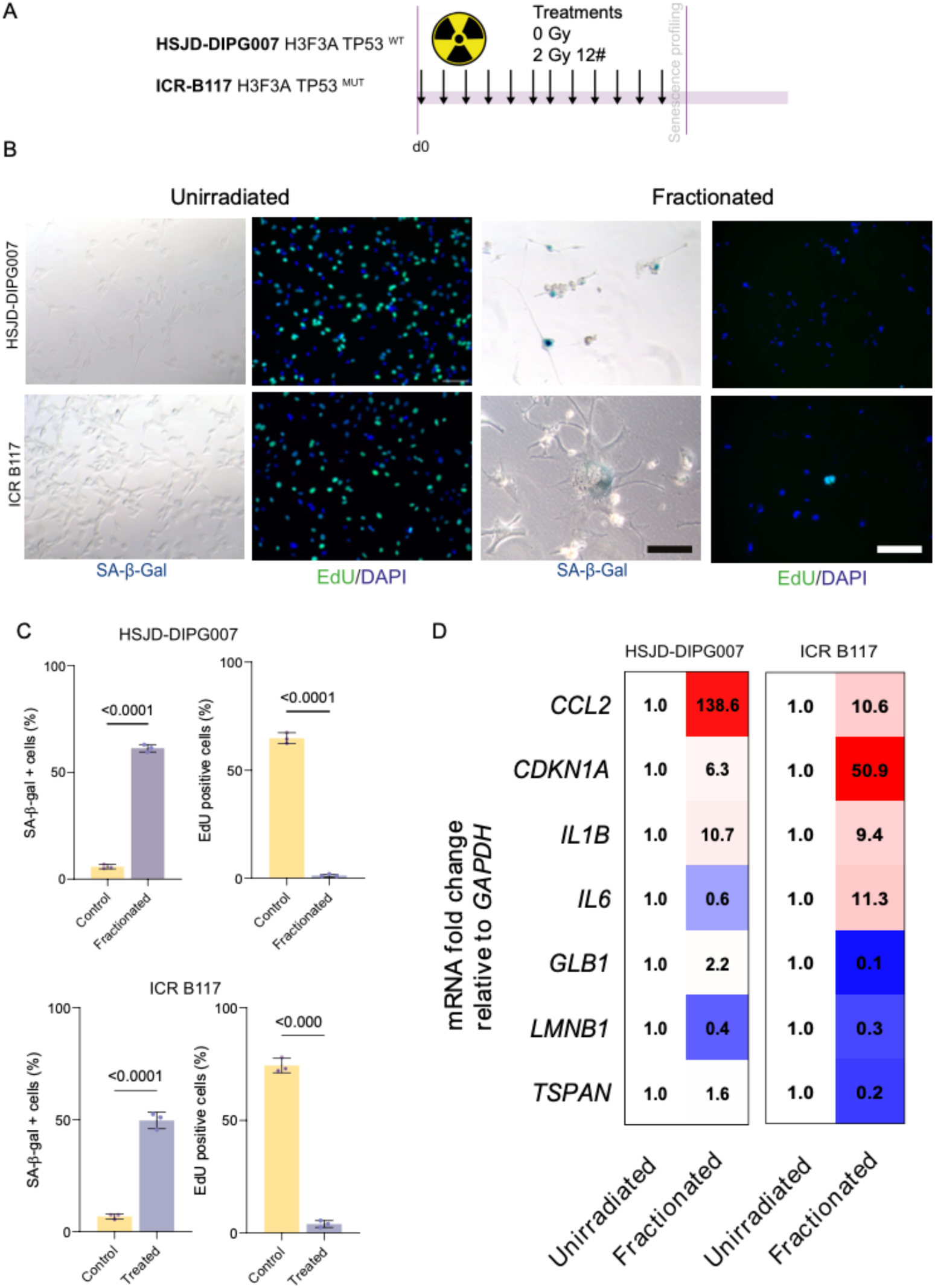

**Suppl. Figure 2:**
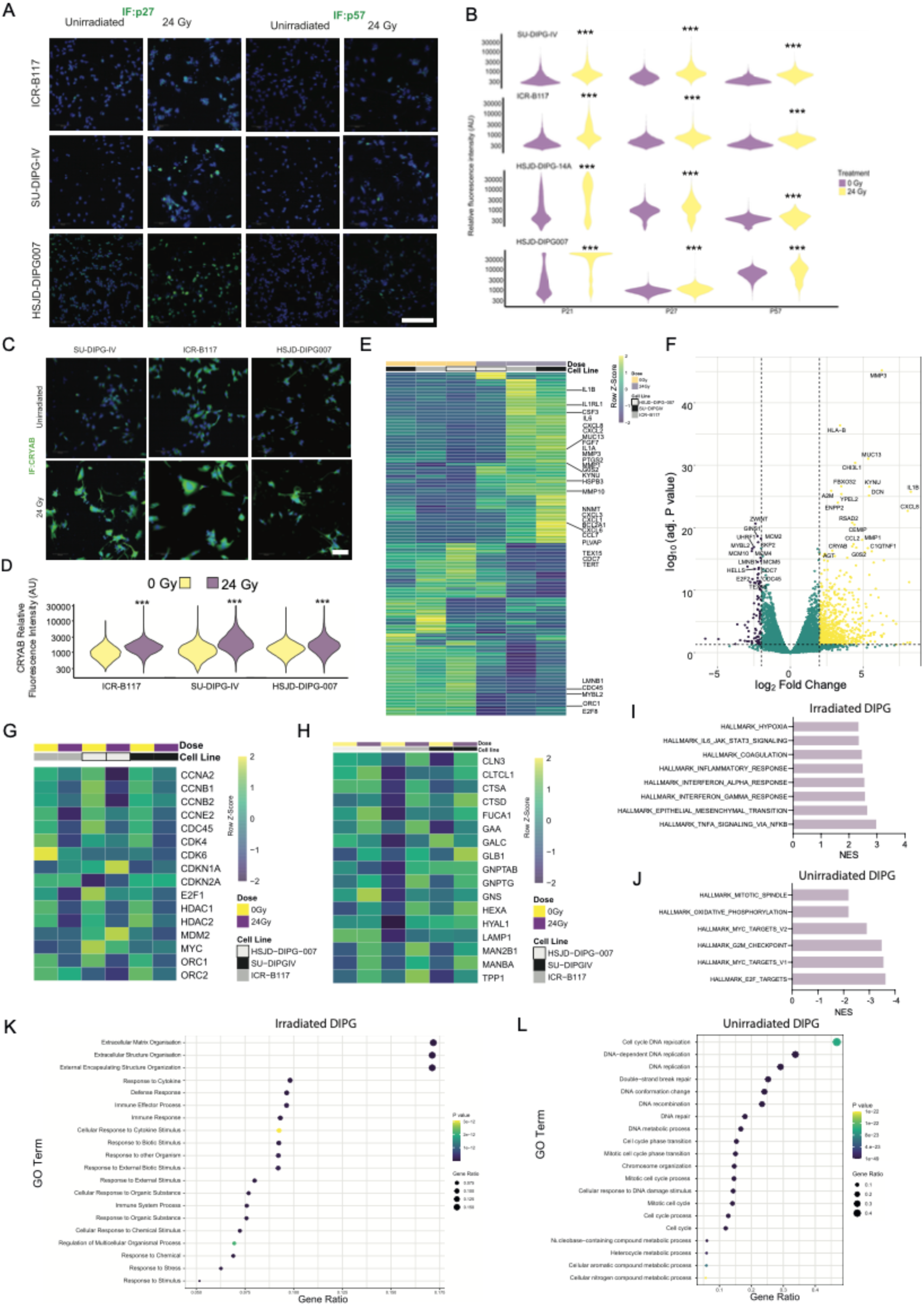

**Suppl. Figure 3:**
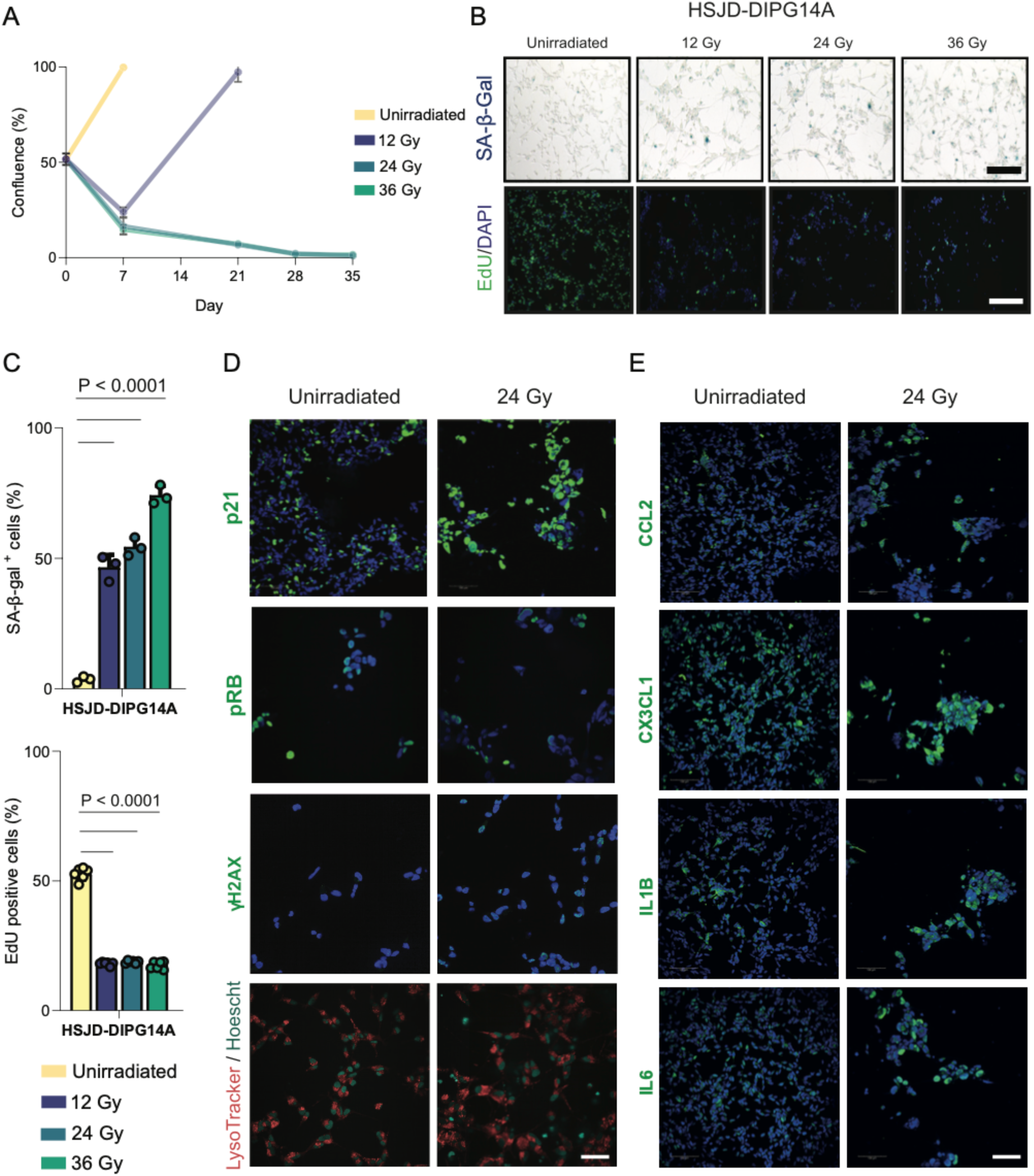

**Suppl. Figure 4:**
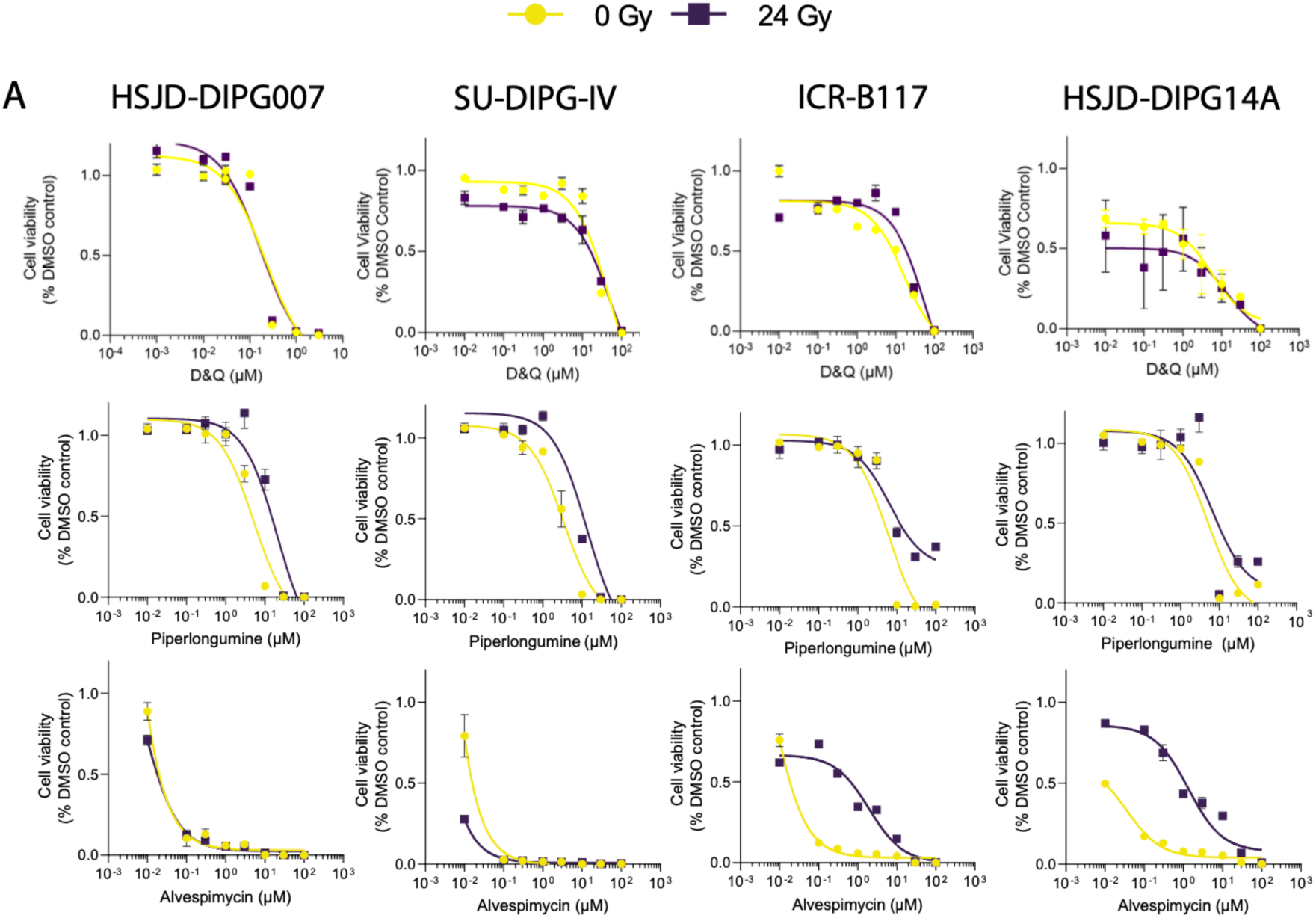

**Suppl. Figure 5:**
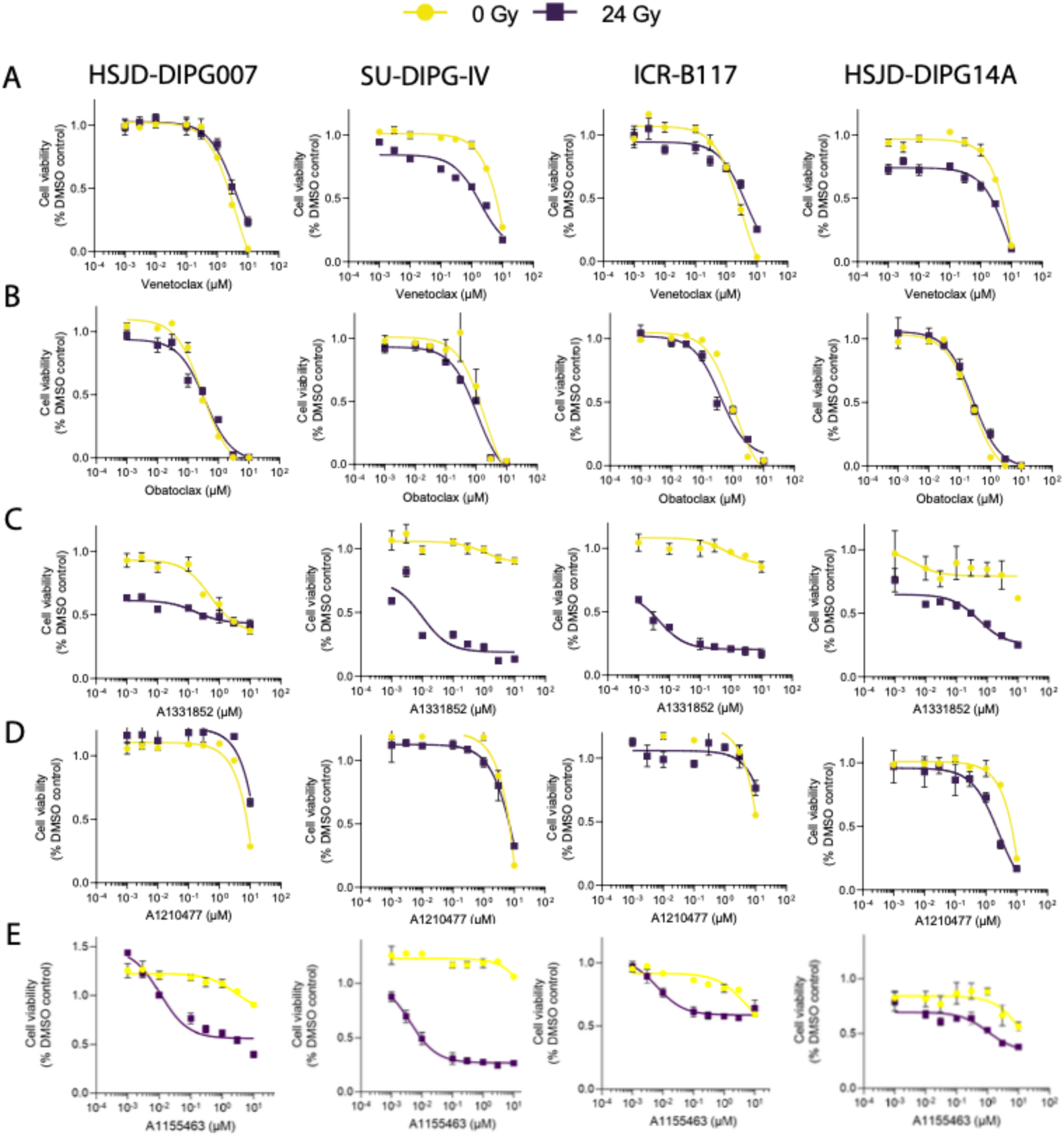

**Suppl. Figure 6:**
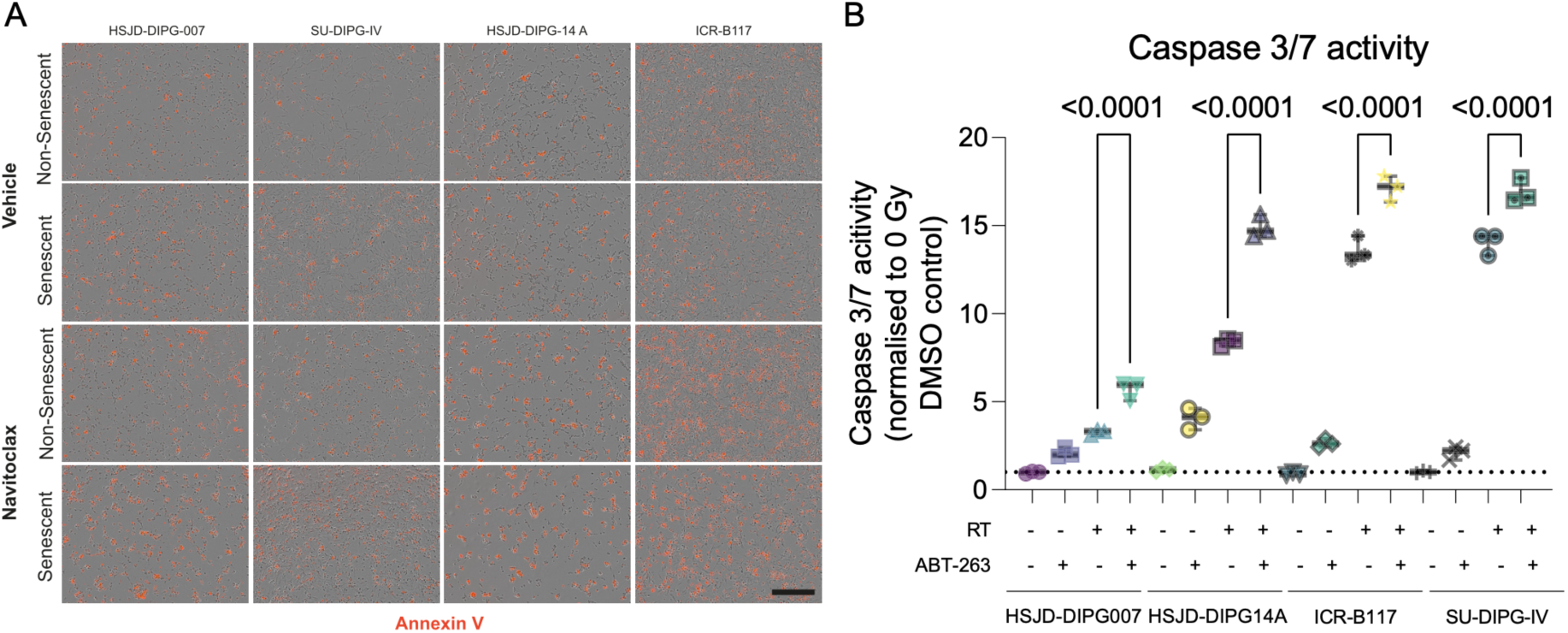

**Suppl. Figure 7:**
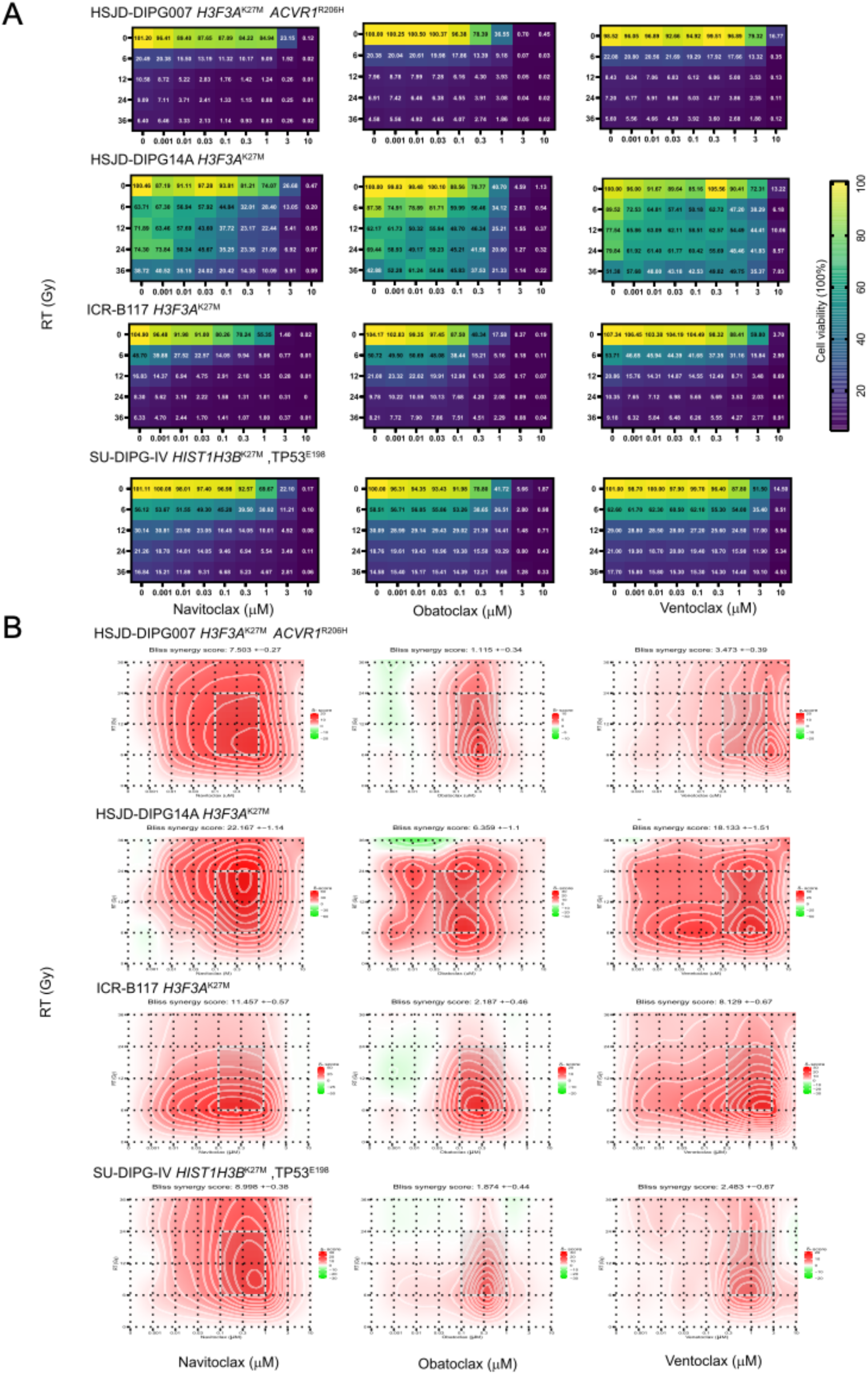

**Suppl. Figure 8:**
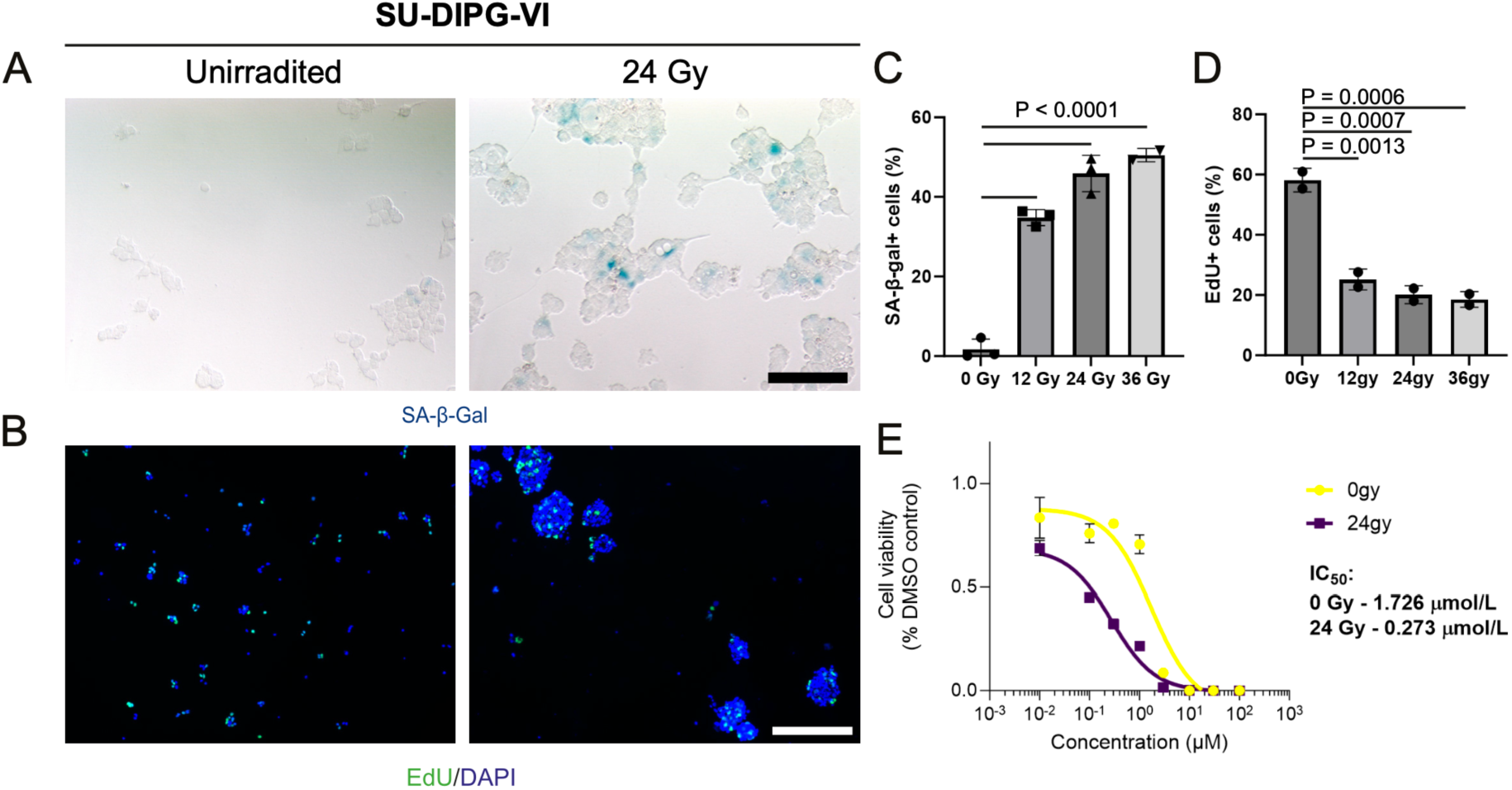

**Suppl. Figure 9:**
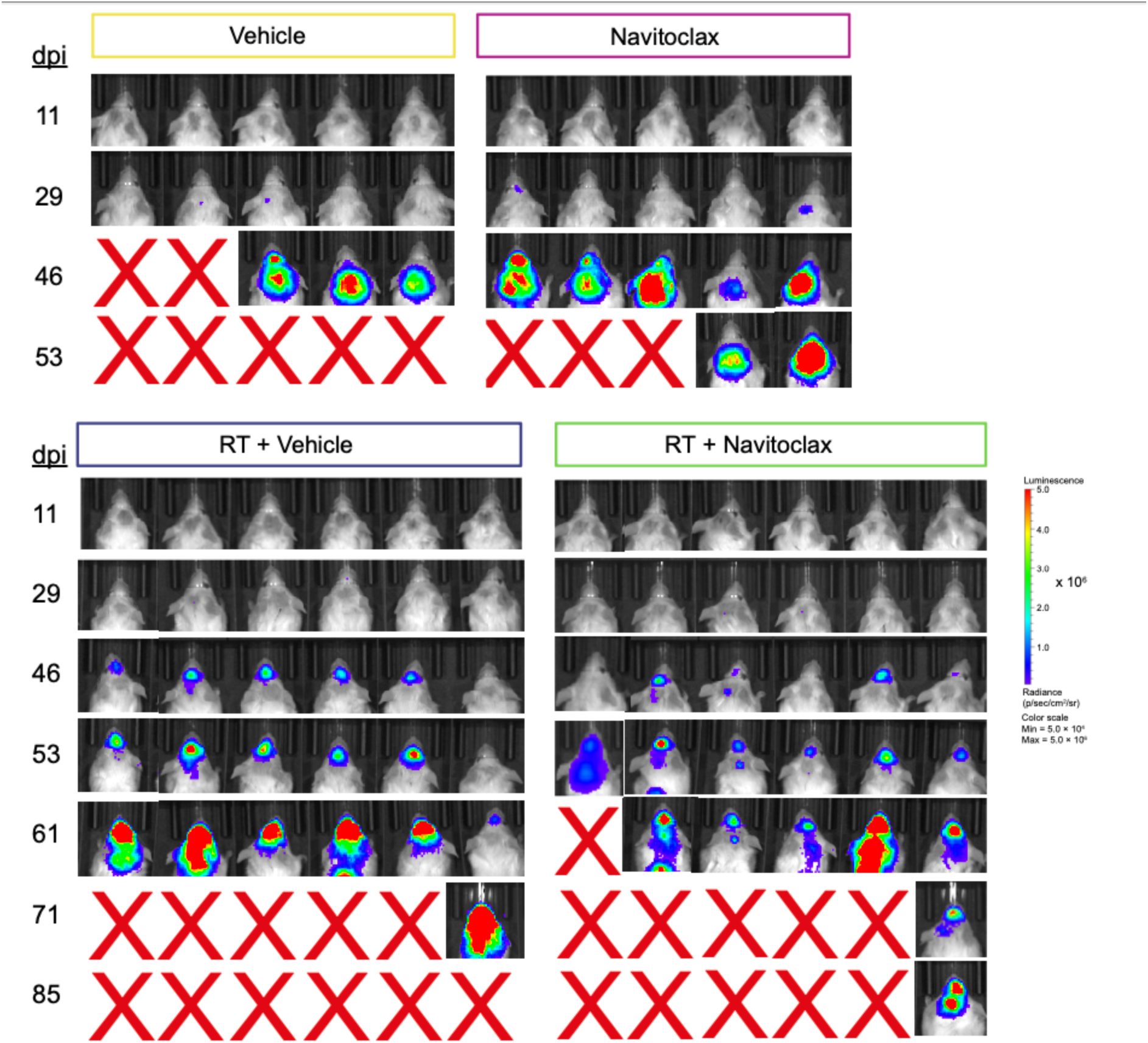

**Supplementary Table 1.** Cell lines used in this study.

| **C e l l**<br>**l i n e**<br>**I D** | **Clinical**<br>**diagnosis** | **Location** | **Age**<br>**(yrs)** | **Sex** | **Histology** | **W H**<br>**O** | **Source** | **Survival**<br>**(month)** | **Outcome** | **Histone**<br>**H3** | **p53** | **ACVR**<br>**1** |
| --- | --- | --- | --- | --- | --- | --- | --- | --- | --- | --- | --- | --- |
| **H S J**<br>**D -**<br>**DIPG**<br>**-007** | DIPG | Pons | 9 | Male | DIPG | IV | Biopsy | 0.9 | Died | H3F3A | WT | R 2 0 6<br>H |
| **H S J**<br>**D -**<br>**DIPG**<br>**-14A** | DIPG | Pons | 8 | Female | DIPG | - | Biopsy | - | Died | H3F3A | WT | WT |
| **I C R -**<br>**B117** | DIPG | Pons | 7 | Male | DIPG | IV | Biopsy | - | Died | H3F3A | p.R273<br>H | WT |
| **S U -**<br>**DIPG**<br>**-IV** | DIPG | Pons | 2 | Female | DIPG | IV | Autopsy | 8.0 | Died | HIST1H3<br>B | p.R273<br>H | G 3 2 8<br>V |
| **S U -**<br>**DIPG**<br>**-VI** | DIPG | Pons | 7 | Female | DIPG | III | Autopsy | 6.0 | Died | H3F3A | p.E198 | WT |

**Supplementary Table 2.** BH3 mimetics used in this study.

| Compound | Inhibitory profile<br>(K<sub>i</sub> cell free assays) |  |  |  | Clinical<br>t r i a l<br>status | E x a m p l e<br>clinical trial | A c h i e v a b l e<br>non-toxic C<sub>max</sub><br>(nM) |
| --- | --- | --- | --- | --- | --- | --- | --- |
|  | Bcl-2 | B c l - | Bcl-w | Mcl-1 |  |  |  |
| **Venetoclax** | <0.01 | 48 | 245 | >440 | M H R A<br>approved | NCT04401748 | 2533.4 ± 1612 |
| **Navitoclax** | <1 | <0.05 | <1 | 550<br>(+/-40) | Phase III | NCT02591095 | 6607 ± 3262 |
| **Obatoclax** | 0.2 | 1-7 | 1-7 | 1-7 | Phase I | NCT00600964 | 3-15.7ng/ml |
| **A-1331852** | 6 | <0.01 | 4 | 142 | Preclinical | n.a | n.a. |
| **A-1155463** | 74 | <0.01 | 8 | >444 | Preclinical | n.a | n.a |
| **A-1210477** | 132 | >660 | 2,280 | <0.5 | preclinical | n.a | n.a |

**Supplementary Table 3.** Primary antibodies used in this study.

| Antibody<br>Application | Target | H o s t<br>Species | Clonality | Clone | Company | Catalogue<br>Number | Dilution | A n t i g e n<br>Retrieval<br>Method |
| --- | --- | --- | --- | --- | --- | --- | --- | --- |
| **Primary** | H u m a n<br>n u c l e i<br>antigen | Mouse | Monoclonal | N/A | Millipore | MAB1281 | 1:500 | Tris- EDTA<br>pH9.0 |
| **Primary** | CX3CL1 | Rabbit | Monoclonal | N/A | R&D | AF472 | 1:1000 | Tris- EDTA<br>pH9.0 |
| **Primary** | CRYAB | Mouse | Monoclonal | 1B6.1-3<br>G4 | Abcam | R ab13496 | 1:300 | Tris- EDTA<br>pH9.0 |
| **Primary** | Ki67 | Rabbit | Monoclonal | SP6 | Abcam | ab16667 | 1:100 | Tris- EDTA<br>pH9.0 |
| **Primary** | P21 | Rabbit | Polyclonal | M-19 | C e l l<br>Signaling<br>Technology | #5487 | 1:400 | Tris- EDTA<br>pH9.0 |
| **Primary** | P57 | Mouse | Monoclonal | 10B1 | C e l l<br>Signaling<br>Technology | #2557 | 1:200 | Tris- EDTA<br>pH9.0 |
| **Primary** | P16 | Mouse | Monoclonal | N/A | Santa Cruz | Sc-56330 | 1:100 | Tris- EDTA<br>pH9.0 |
| **Primary** | P27 | Mouse | Monoclonal | VE1 | C e l l<br>Signaling<br>Technology | E19290 | 1:50 | Tris- EDTA<br>pH9.0 |
| **Primary** | p R B<br>(Ser807/81<br>1) | Rabbit | Polyclonal | N/A | C e l l<br>Signaling<br>Technology | 9308 | 1:250 | Tris- EDTA<br>pH9.0 |
| **Primary** | CCL2 | Rabbit | Monoclonal | N/A | R&D | MAB679R | 1:1000 | Tris- EDTA<br>pH9.0 |
| **Primary** | IL1B | Rabbit | Monoclonal | N/A | R&D | MAB201 | 1:200 | Tris- EDTA<br>pH9.0 |
| **Primary** | IL6 | Rabbit | Monoclonal | N/A | R&D | AF-206-NA | 1:1000 | Tris- EDTA<br>pH9.0 |
| **Primary** | H3K27M | Rabbit | Polyclonal | N/A | Abcam | Ab19061 | 1:500 | Tris- EDTA<br>pH9.0 |

**Supplementary Table 4.** Secondary antibodies used in this study.

| **Antibody Application** | **Target Species** | **Host Species** | **Clonality** | **Company** | **Catalogue Number** | **Dilution** | **Conjugate** |
| --- | --- | --- | --- | --- | --- | --- | --- |
| **Secondary** | Mouse | Goat | Polyclonal | Thermo Fisher | A11001 | 1:250 | Alexa Fluor® 488 |
| **Secondary** | Mouse | Goat | Polyclonal | Thermo Fisher | A11004 | 1:250 | Alexa Fluor® 568 |
| **Secondary** | Rabbit | Goat | Polyclonal | Thermo Fisher | A11008 | 1:250 | Alexa Fluor® 488 |
| **Secondary** | Rabbit | Goat | Polyclonal | Thermo Fisher | A11036 | 1:250 | Alexa Fluor® 568 |
| **Secondary** | Goat | Chicken | Polyclonal | Thermo<br>Fisher | A21467 | 1:250 | Alexa Fluor®<br>488 |
| **Secondary** | Mouse | Goat | Polyclonal | Dako | E0433 | 1:250 | Biotin |
| **Secondary** | Rabbit | Goat | Polyclonal | Dako | E0432 | 1:250 | Biotin |
| **Secondary** | Mouse | Goat | Polyclonal | Dako | P0447 | 1:250 | Horseradish<br>peroxidase<br>(HRP) |

